# Predicting potentially permissive substitutions that improve the fitness of A(H1N1)pdm09 viruses bearing the H275Y NA substitution

**DOI:** 10.1101/2021.03.21.436293

**Authors:** Rubaiyea Farrukee, Vithiagaran Gunalan, Sebastian Maurer-Stroh, Patrick C. Reading, Aeron C. Hurt

## Abstract

Oseltamivir-resistant influenza viruses arise due to amino-acid mutations in key residues, but these changes often reduce their replicative and transmission fitness. Widespread oseltamivir-resistance has not yet been observed in A(H1N1)pdm09 viruses. However, it is known that permissive mutations in the neuraminidase (NA) of former seasonal A(H1N1) viruses from 2007-2009 buffered the detrimental effect of the NA H275Y mutation, resulting in fit oseltamivir-resistant viruses that circulated widely. This study explored two approaches to predict permissive mutations that may enable a fit H275Y A(H1N1)pdm09 variant to arise.

A computational approach used phylogenetic and *in silico* protein stability analyses to predict potentially permissive mutations, which were then evaluated by *in vitro* NA enzyme activity and expression analysis, followed by *in vitro* replication. The second approach involved the generation of a virus library which encompassed all possible individual 2.9 × 10^4^ codon mutations in the NA whilst keeping H275Y fixed. To select for variant viruses with the greatest fitness, the virus library was serially passaged in ferrets (via contact and aerosol transmission) and resultant viruses were deep sequenced.

The computational approach predicted three NA permissive mutations, and even though they only offset the *in vitro* impact of H275Y on NA enzyme expression by 10%, they could restore replication fitness of the H275Y variant in A549 cells. In our experimental approach, a diverse virus library (97% of 8911 possible single amino-acid substitutions were sampled) was successfully transmitted through ferrets, and sequence analysis of resulting virus pools in nasal washes identified three mutations that improved virus transmissibility. Of these, one NA mutation, I188T, has been increasing in frequency since 2017 and is now present in 90% of all circulating A(H1N1)pdm09 viruses.

Overall, this study provides valuable insights into the evolution of the influenza NA protein and identified several mutations that may potentially facilitate the emergence of a fit H275Y A(H1N1)pdm09 variant.

## 1. Introduction

Oseltamivir is a neuraminidase inhibitor (NAI) which is widely used and prescribed for the treatment of influenza, and is often stockpiled for pandemic purposes [1-5]. This drug was designed to target the conserved active site of the influenza virus neuraminidase (NA) glycoprotein and inhibit its enzymatic function, hence reducing the capacity of virus to release from infected host cells [6, 7]. However, amino acid substitutions that arise in the drug binding region of the NA glycoprotein can reduce virus susceptibility to oseltamivir [8, 9]. For example, the H275Y amino acid substitution that is commonly reported in the NA of influenza A(H1N1) viruses [1-4,10, 11] prevents the conformational change of the E276 amino acid which normally creates a hydrophobic pocket necessary for oseltamivir binding, and leads to reduced oseltamivir susceptibility [12-15]. Therefore, the emergence of viruses bearing this substitution is of particular concern.

Prior to 2008, the prevalence of the H275Y NA substitution in former seasonal A(H1N1) viruses was generally low (<1%) [11, 16-19]. Earlier *in vitro* and *in vivo* studies, often performed with older laboratory viruses such as A/WSN/33, A/New Caledonia/20/99 and A/Texas/36/91, showed that that variants with the H275Y NA substitution had reduced NA enzyme function, and reduced replication and transmission capabilities compared to wild-type viruses [12, 20-24]. Given the observed fitness loss it was assumed that this substitution was unlikely to circulate widely amongst the community. However, in 2008, an H275Y variant emerged in the A/Brisbane/59/2007-like A(H1N1) virus background, that was able to outcompete all circulating wild-type strains and reach nearly 100% frequency [25-29]. Fortunately seasonal A(H1N1) viruses bearing the H275Y substitution were replaced by swine origin A(H1N1)pdm09 viruses in 2009/2010 and these new viruses retained sensitivity to oseltamivir [30-32]. However, the rapid emergence of A/Brisbane/59/2007-like A(H1N1) viruses with the H275Y NA substitution highlighted the potential for H275Y variants to be fit and transmissible, and demonstrated the need to closely monitor the evolution of the NA glycoprotein of A(H1N1)pdm09 viruses.

To gain insights into the factors facilitating the emergence of a transmissible A(H1N1) variant with H275Y NA substitution, analyses have been performed to compare the effect of the H275Y substitution in the permissive A/Brisbane/59/2007 virus background with that of older virus strains [33].These *in vitro* and *in vivo* replication and transmission studies showed that the H275Y NA substitution did not impact the fitness of the A/Brisbane/2007-like viruses to the same extent it did to older virus strains such as A/WSN/33 [34-36]. Subsequent analyses demonstrated that due to the acquisition of certain substitutions (R222Q, V234M and D344N), the NA from A/Brisbane-like viruses had different enzymatic properties compared to NAs from earlier seasonal A(H1N1) viruses and, were able to restore the deficits in NA enzyme function due to H275Y [36-41]. Substitutions in the HA (T82K, K141E and R189K) were also found to play a role in restoring the fitness of A/Brisbane-like viruses [42]. These studies highlighted that virus evolution can lead to incorporation of permissive substitutions in viral NA, that can in turn facilitate the emergence and spread of H275Y in circulating viruses.

Currently, the prevalence of the H275Y NA substitution in circulating A(H1N1)pdm09 viruses is low (<1%) [1-5,43]. To date, experimental studies assessing viral fitness have shown mixed results, with some reporting comparable fitness between wild-type virus and H275Y variants [44-48], while others reporting impaired fitness of H275Y variants [49-51]. However, clusters of A(H1N1)pdm09 variants with the H275Y have been reported in community settings, notably in Australia in 2011 [52] and in Japan in 2014 [53]. Detailed analysis of the viruses from the 2011 Australian cluster demonstrated that the A(H1N1)pdm09 viruses had acquired permissive NA substitutions V241I and N369K, which partially restored the fitness deficit due to H275Y [54, 55]. Interestingly, a previous study had utilised computational analyses to predict that the N369K could be potentially permissive for H275Y in A(H1N1)pm09 viruses [56]. The V241I and N369K substitution are now present in all circulating viruses [54], but given the low prevalence of H275Y in currently circulating viruses, further permissive substitutions are likely needed for H275Y to become widespread. This is supported by a recent study, which utilised A(H1N1)pdm09 viruses from 2016 to demonstrate that variants with the H275Y substitution still showed reduced fitness compared to the corresponding wild-type virus, although not to the same extent as the H275Y substitution in a 2009 A(H1N1)pdm09 virus [57].

A key lesson from the widespread circulation of a fit H275Y variant in 2008 was that virus evolution can lead to substitutions in viral NA, which allows the virus to become permissive for the H275Y substitution. Since this phenomenon is possible again in the newer swine-origin A(H1N1)pmd09 viruses, our aim is to identify possible substitutions that may emerge in the NA of this virus to facilitate the emergence and spread of H275Y variants. To do end, we have used two different approaches to predict possible permissive NA substitutions in influenza A(H1N1)pdm09 viruses, which might offset the fitness loss due to H275Y. In our first approach, we have used a computational analysis to predict possible candidates for permissive NA substitutions, followed by *in silico* calculations to ascertain their impact on protein stability. Our second approach involved the generation of a virus library which was designed to contain every possible single amino acid substitution in the viral NA, while keeping the H275Y fixed. The virus library was then used to infect ferrets via serial transmission to select for variants with high fitness, and thereby identify candidates for permissive substitutions. A selection of the candidate NA substitutions identified using either the computational or experimental approach described above were analysed further, to determine their effect on NA cell-surface expression and activity, and virus replication. The data obtained from these experiments has allowed us to propose several candidate substitutions that may make a A(H1N1)pdm09 virus potentially permissive for H275Y in the future.

## 2. Materials and Methods

### 2.1 Computational approach to predict permissive substitutions

#### 2.1.1 Bioinformatics analysis

The computational approach included analysis of N1 protein sequences from human A(H1N1)pdm09 viruses available from Global Initiative on Sharing All Influenza Data website (http://www.gisaid.org) and the influenza virus resource at the National Centre for Biotechnology Information and followed by *in silico* protein stability calculations. Briefly, substitutions that could potentially be permissive for H275Y were selected through the following criteria: a) substitutions that have co-occurred with H275Y in A(H1N1)pdm09 viruses, and b) were present in a minimum of 10 sequences. A selection of 25 substitutions were identified which were then grouped into sets of four, yielding 12,650 possible combinations representing the serial accumulation of each substitution in all combinations, prior to the acquisition of H275Y. The FoldX program [58] was then used to calculate the effect of these substitution sets on the Gibbs free energy (energy of unfolding, ΔG, kcal/mol) of a representative three-dimensional NA protein structure. The change in free energy (ΔΔG) from wild-type protein was calculated for each set of substitution, and permissive pathways were constructed representing serial addition of each substitution in a set using a custom Perl script. A fitness threshold was selected based on previous studies done with H275Y variants in Newcastle, Australia in 2011 [52] and energy changes were calculated relative to the NA from A/California/07/2009). The background NA structure was derived by homology modelling with Modeller [59] using A/California/04/2009 (PDB ID: 3NSS) as a template.

### 2.2 Overview of Experimental approach for selecting functional variants

To assess the impact of all possible amino acid substitutions on viral fitness, a virus library was produced that expressed all possible individual codon mutations (2.9 × 10^4^) in the NA whilst keeping H275Y fixed. The virus library was then passaged through ferrets by serial transmission (n = 4 independent lines of transmission) to select for functional variants. Deep sequencing was performed on the virus library and on ferret nasal washes on selected days, to ensure the completeness of the library and, to identify which amino acids were under positive selection pressure in the presence of H275Y.

#### 2.2.1 Generation of Virus library

The first step for creating the virus library involved codon-based mutagenesis, which was used to generate three independent NA plasmid libraries (i, ii, iii) as has been previously described for influenza A virus HA and NP genes [60-62]. The template NA for the library preparation was from an A(H1N1)pdm09 virus, A/South Australia/16/2017, with the H275Y substitution (H275Y-NA) introduced by site-directed mutagenesis using the GeneArt site directed mutagenesis kit (Invitrogen, USA). The A/South Australia/16/2017 virus isolate (herein referred to as the SA16-WT virus), was submitted to the WHO Collaborating Centre for Reference and Research in Melbourne, Australia as part of the WHO GISRS surveillance programme.

Each replicate of the NA plasmid library (i, ii and iii) was then used to generate a virus library by reverse genetics (i, ii and iii), as has been described previously [61, 63, 64]. Briefly, co-cultures of 293T and MDCK-SIAT1-TMPRRS2 [65] cells were transfected with the pHW2000 plasmid containing the seven genes from the SA16-WT virus, and with either NA plasmid library i, ii or iii. Overall, four viral rescues were performed: three with the NA plasmid library replicates (resulting in virus libraries i, ii, and iii) and one control rescue with just the H275Y-NA plasmid (resulting in the SA16-H275Y virus). In order to control for loss of viral diversity due to bottlenecks introduced during reverse genetics, each rescue was done in replicates of six and the supernatants from the replicates were then pooled together to create each virus library. Of note, as a further precaution, the MP gene for all viruses were modified to revert the S31N mutation, so viruses retained sensitivity to adamantanes. Titres of infectious virus in the virus library preparation were determined using a TCID_50_ assay [66].

A more detailed description of the library preparation is available in Supplementary text S1.

#### 2.2.2 Selection for functional variants in the ferret model

##### Ethics statement

Experiments using ferrets were conducted with approval from the Melbourne University Animal Ethics Committee (project license number 1714278.) in strict accordance with the Australian Government, National Health and Medical Research Council Australian code of practice for the care and use of animals for scientific purposes (8^th^ edition). Animal studies were conducted at the Bio Resources Facility located at the Peter Doherty Institute for Infection and Immunity, Melbourne.

##### Ferrets

Outbred adult male and female ferrets older than 6-months and weighing 608–1769g were used. Prior to inclusion in experiments, serum samples were collected and tested by hemagglutination inhibition assay [67] against reference strains of influenza A and B viruses to ensure seronegativity against currently circulating influenza subtypes and lineages. Ferrets were housed individually in high efficiency particulate air filtered cages with *ab libitum* access to food, water and enrichment equipment throughout the experimental period. Ferrets were randomly allocated to experimental groups.

The three virus libraries (i, ii and iii) generated by reverse genetics were pooled into a single virus library to increase the likelihood that all possible substitutions were comprehensively sampled in the final virus library. This final library was subsequently passaged through ferrets in 4 independent lines of transmission to select for variant viruses with the greatest fitness.

Four ferrets were experimentally inoculated with 500 µL containing 10^4.7^ TCID_50_ of pooled virus library (day 0), as previously described [68]. One ferret was experimentally inoculated with the SA16-H275Y virus as a control. Each, experimentally infected ferret was then co-housed with a naïve contact recipient (direct contact 1) 24 hours post-inoculation. Nasal washes were performed daily on direct contact 1 ferrets and nasal wash samples were analysed for infection by qPCR [54]. On the first day that nasal wash samples from direct contact 1 ferrets were qPCR positive for influenza virus, the animal was removed from the cage, and co-housed with a second naïve recipient (direct contact 2). Similarly, nasal wash samples from direct contact 2 were monitored for influenza virus. On the first day that nasal wash samples from direct contact 2 ferrets were qPCR positive for influenza virus, these animals were placed in aerosol cages, adjacent to a third set of naïve recipients (aerosol contacts). Due to limited animal numbers, the SA16-H275Y virus was only passaged once through ferrets (Experimentally infected animals to Direct Contact 1).

In the experiments described, ferrets were nasal washed every day and weight and body temperatures were collected as previously described [69]. Experimentally infected ferrets were euthanized on day 4 of the experiment, and all other animals were euthanized on day 14 of the experiment. Viral titres in nasal wash samples were determined by qPCR [54] and TCID_50_ assay [66].

Figure 1 presents an overall schematic for the serial transmission experiments in the ferret model.

**Figure 1:**
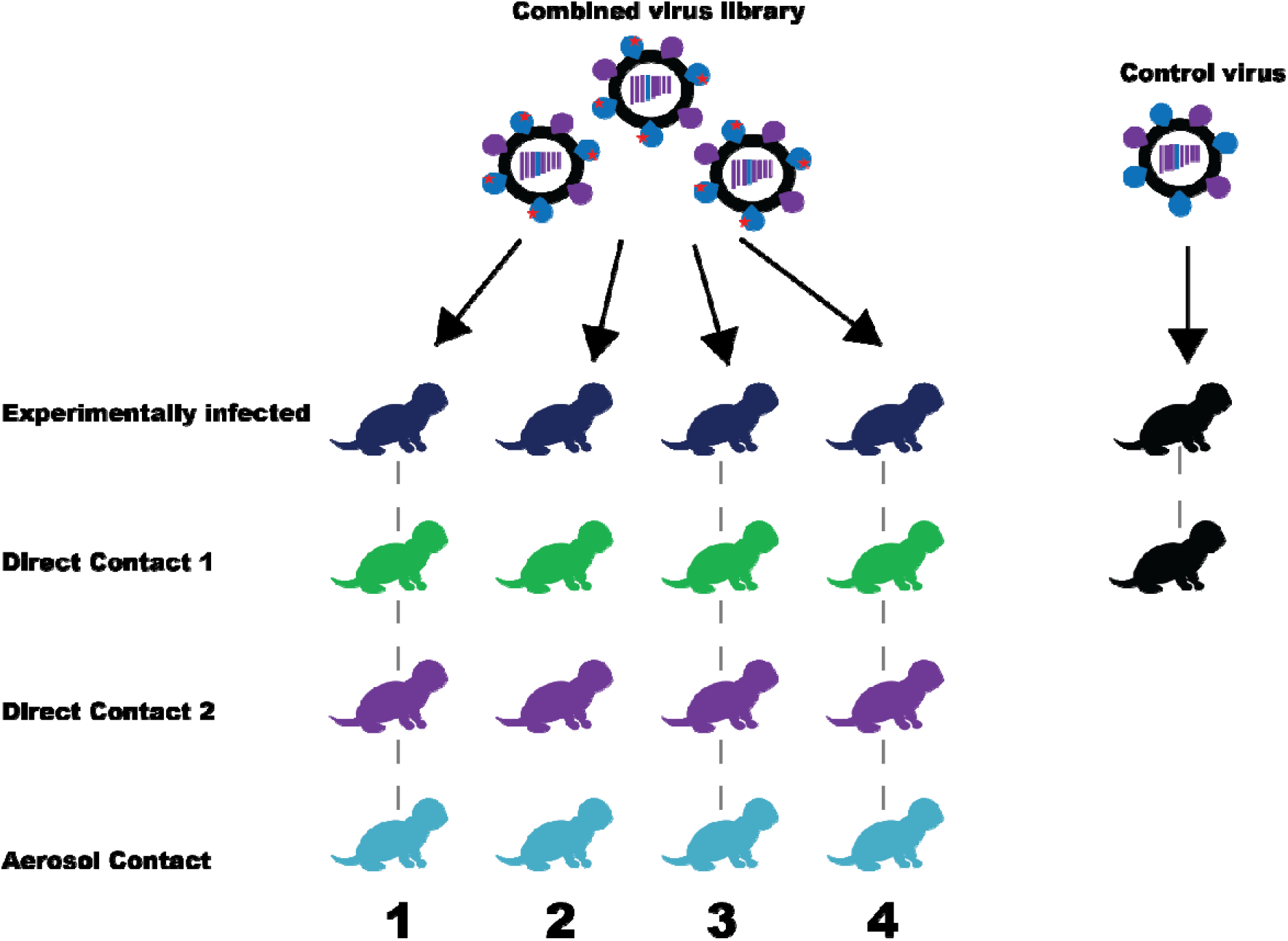
Schematic of the transmission model used to select for fit H275Y variants in the ferret model of influenza infection. Codon-based mutagenesis and reverse genetics was used to generate virus libraries in triplicates, such that it contained viruses with all possible codon mutations in the A/South Australia/16/2017-NA with the exception of H275Y substitution. The virus libraries were pooled together to increase the likelihood that all codon mutations were represented, and the combined library was passaged through ferrets via serial transmission (n = 4 independent lines of transmission) and nasal wash samples were collected and analysed to determine if any variant had been selected from the virus library via passage through ferrets. As a control, the A/South Australia/16/2017-H275Y virus, generated by reverse genetics, was passaged through ferrets once, to determine the background mutation frequency.

#### 2.2.3 Deep sequencing analysis of virus library and ferret nasal washes

The plasmid libraries (i, ii and iii) and virus libraries (i, ii and iii) were deep sequenced, alongside the H275Y-NA plasmid (control for PCR error rate) and the SA16-H275Y virus (control for reverse genetics). A single nasal wash sample was picked from each ferret in the transmission chain for deep sequencing (sample selection is denoted in Figure 4). Factors taken into consideration when selecting nasal wash samples for deep sequencing were (i) selection of time points as late as possible during infection to allow time for within-host selection of variants from the viral mixtures, and (ii) that the RNA quantity and quality was sufficient for deep sequencing and accurate variant calling [70].

Viral RNA from ferret nasal wash samples was quantified using qPCR with primers that detect the M gene of influenza A viruses, provided by the US Centers for Disease Control and Prevention, Atlanta, USA.

Viral RNA was extracted from virus library and ferret nasal wash samples using the QIAamp® Viral RNA mini kit (Qiagen, Germany). Next generation sequencing was carried out twice on the nasal washes: a) on the NA gene only to get a high degree of coverage for analysis b) for the full genome of the virus to track changes in the internal genes. For the NA gene, cDNA synthesis was carried out using NA gene-specific primers (supplementary text S1), and the SuperScript III First-Strand Synthesis System (Invitrogen, USA). The NA gene was amplified from the cDNA and plasmids using gene-specific primers and the Platinum™ *Taq* DNA Polymerase High Fidelity kit (Invitrogen, USA) and sent for sequencing. The full genome sequencing was done after amplification of all genes using primers previously described [71]. Sequencing of amplified PCR products were done at the Australian Genome Research Facility, on the HiSeq 2500 platform (2x 150 PE reads, 15 million reads per sample).

#### 2.2.4 Analysis of deep sequencing data and bioinformatics

The NA genes from the plasmid and virus libraries were deep sequenced alongside their respective controls to confirm that all single amino acid substitutions were represented in each library. The mapmuts pipeline (http://jbloom.github.io/mapmuts/) was used to generate codon counts for each site. Codon identities were called only in overlapping regions of the paired-end reads, where both reads concurred. This was done to reduce the sequencing error rate, as the same sequencing error is unlikely to occur in both reads. The dms_tools2 and mapmuts pipelines were then used to confirm the completeness of the libraries. The pipelines were also used to map overlapping fastq reads from ferret nasal washes to template NA [37,60-62,65].

For ferret nasal wash samples, fastq reads were also mapped to the influenza genome using Bowtie2 v2.2.5 (-very-sensitive-local) (http://bowtiebio.sourceforge.net/index.shtml). SAM tools v1.7 was used to process sequence alignments and generate pileup files. The pileup files were then used to scan for minorities using Varscan [75] with a minimum variant calling threshold set at 1%. The nucleotide diversity and ratio of synonymous to non-synonymous mutations in ferret nasal wash samples was calculating by measuring π and πS/πN using the SNPgenie software [76]. The nucleotide mutation frequencies in donor:recipient pairs from the eight contact transmission pairs and for aerosol transmission pairs were also used to estimate transmission bottleneck sizes using the beta-binomial sampling method developed by Leonard *et al* [77]. This statistical method takes the stochastic dynamics of viral replication in recipients into account and further considers variant calling thresholds. For our analysis, a minimum variant calling threshold of 1% was utilised to estimate bottleneck size to include a greater number of sites, as was done by Poon *et al*. in a human household transmission study [78]. A more conservative estimate of the bottleneck size was also calculated, using a minimum variant calling threshold of 3% similar to Leonard *et al*. [77].

#### 2.2.4 Sequence Availability

Illumina sequencing data are available at the Sequence Read Archive (SRA) (Accession PRJNA561026, https://www.ncbi.nlm.nih.gov/sra/PRJNA561026, last accessed 27^th^ January, 2021).

### 2.3 Evaluation of candidate permissive substitutions on viral fitness

#### 2.3.1 NA cell surface expression and activity assay

To gain some insights regarding the impact of each candidate substitution identified by computational or experimental approaches described above, we investigated the effect of these substitutions on NA cell-surface expression and NA activity. For these experiments, the H275Y-NA gene was incorporated into an expression plasmid with a V5 epitope tag and appropriate substitutions were introduced by site-directed mutagenesis. Measurement of cell-surface NA expression and activity was performed by transfecting 293T cells with the expression plasmid as has been described in our previous studies [37,54,56,79,80]. Three independent experiments were performed to assess NA expression and activity, where each variant NA was tested in triplicate. GraphPad Prism v.6 was used for statistical analysis of between group comparison differences using an unpaired Student ‘s two-tailed *t*-test.

#### 2.3.2 Virus replication in A549 cells

The most promising candidate substitutions from the previous analyses were incorporated into the SA16-H275Y virus by site-directed mutagenesis and reverse genetics. The replication kinetics of the SA16-H275Y virus, the SA16-WT virus (generated by reverse genetics instead of using isolate to maintain consistency), and the SA16-H275Y viruses with candidate substitutions was then evaluated in A549 cells (lung carcinoma cell lines), infected with an MOI of 0.1. The multi-cycle replication kinetics for each virus was performed in triplicates and viral titres were determined at 2, 24, 48 and 72 hours post-infection.

## 3 Results

### 3.1 Computational Approach proposed three candidate substitutions which worked synergistically to improve viral fitness

Bioinformatics analyses were performed to identify substitutions that may be permissive for the H275Y substitution in the N1 NA background. This analysis looked at substitutions that co-occurred with H275Y and were found to occur in at least 10 different viruses. There were 25 such substitutions found which were then used to reconstruct possible permissive pathways *in silico*, and measure impact on protein stability (by calculating change in free energy) (Figure S1). Of these, 15 substitutions were shown to improve protein in stability *in silico* and amongst these, three substitutions were chosen as they were most frequently observed in the reconstructed permissive pathways: S95N (66 pathways), S299A (99 pathways) and S286G (315 pathways) (Figure S1).

The impact of these substitutions in offsetting the fitness loss due to H275Y was then measured experimentally. Firstly, their impact on NA enzyme function was measured individually and in all possible combinations with each other. The results showed that the introduction of the H275Y substitution reduced relative NA activity to 65 ± 9% of the wild-type, and this was not substantially improved by the addition of any of the candidate substitutions (Figure 2A). Introduction of H275Y also reduced NA expression relative to wild-type (48 ± 2%), but a significant improvement in relative NA expression was observed when the S299A substitution was present, with the greatest increase (10%) observed with the combination of S299A+S286G+S95N (Figure 2A). It should be noted however that this increase only partially recovered NA expression relative to wild-type (58 ± 2%), such that expression was still well below 100%.

**Figure 2:**
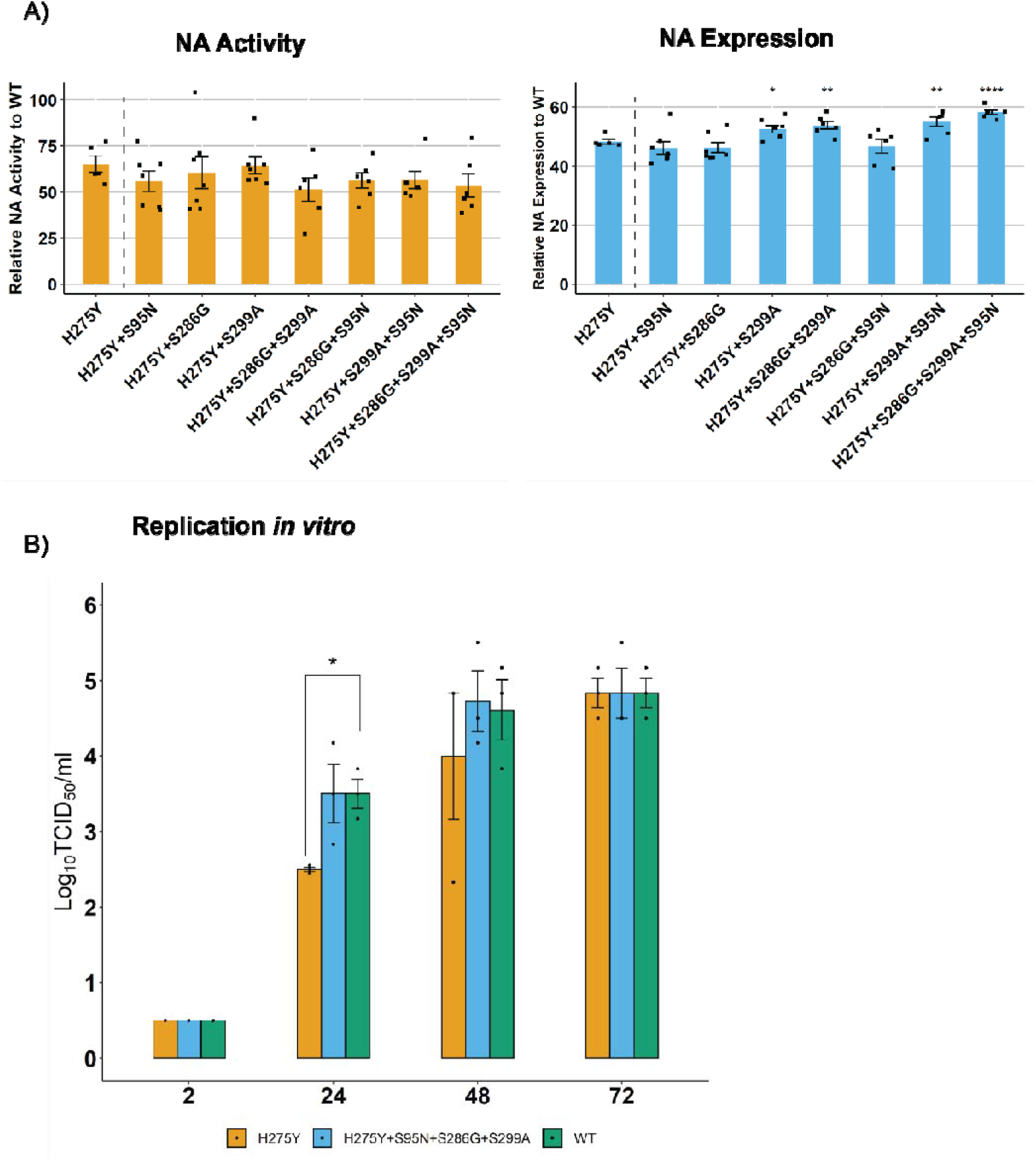
a) The relative NA activity and expression of variant NA glycoproteins with candidate substitutions derived from computational approaches were measured and compared against the H275Y-NA. The NA glycoprotein of the A/South Australia/16/2017 virus was mutated such that it contained the H275Y substitution by itself or in different combinations with candidate permissive substitutions. The proteins were expressed in cells following transfection of 293T cells, and the relative NA activity and expression was calculated as a percentage of wild-type NA protein. Experiments were performed in duplicate on two separate occasions and data are expressed as the mean ± SD. The relative NA activity and expression for the NA proteins containing candidate substitutions were compared against that of the H275Y-NA using a Student ‘s unpaired two-tailed *t*-test. * p<0.05, ** p<0.01 **b)** The replication kinetics of SA16-H275Y, SA16-WT and SA16-H275Y modified with the S95N+S286G+S299A NA substitution was measured in A549 cells infected with an MOI 0.1. The experiment was performed in triplicates and viral titres at each time point were measured using a Student ‘s unpaired two-tailed *t*-test. * p<0.05, ** p<0.01.

Since the combination of S299A+S286G+S95N showed the greatest improvement in enzyme expression, the impact of these substitutions on viral growth kinetics was also tested. Replication kinetics in A549 cells demonstrated delayed growth of SA16-H275Y virus compared to SA16-WT virus, with viral titres reduced at 24 hr (2.5 ± 0.0 Log_10_ TCID_50_/ml vs 3.5 ± 0.2 Log_10_ TCID_50_/ml, p <0.05) and 48 hr (4.0 ± 0.8 Log_10_ TCID_50_/ml vs 4.6 ± 0.4 Log_10_ TCID_50_/ml) post-infection (Figure 2B). Interestingly, the addition of the three substitutions, S299A+S286G+S95N, recovered this delay in virus growth as observed in Figure 2B, suggesting a compensatory/permissive role of these substitutions in regaining loss of viral fitness due to H275Y *in vitro*.

### 3.2 Deep sequence analysis demonstrates that the SA16-H275Y virus library comprehensively sampled all possible amino acid mutations

The experimental approach for identifying permissive substitutions involved creating a virus library by reverse genetics (from a NA plasmid library), and then passaging it through ferrets to select for fit variants. The virus and plasmid libraries were deep sequenced to test for their completeness in sampling all possible amino acid mutations. The reads from these libraries contained at least 10^7^ overlapping paired-end reads aligned to the NA gene and a codon read depth of at least 10^6^ reads per site, which was adequate to sample all mutations present. The per-codon mutation frequency was substantially higher in the plasmid and virus libraries compared to their respective controls (Figure 3A and B). Mutations within the controls consisted of almost entirely single-nucleotide codon changes, as multi-nucleotide changes in the same codon due to sequencing or PCR errors are highly unlikely (Figure 3A). Conversely, the libraries and nasal washes consisted of one-, two-and three-nucleotide changes introduced due to codon mutagenesis. The virus library had a slightly lower rate of per-codon mutation than the plasmid library due to the bottlenecking introduced during reverse genetics, and most of the reduction was in the frequencies of non-synonymous and stop-codon mutations (Figure 3B).

**Figure 3:**
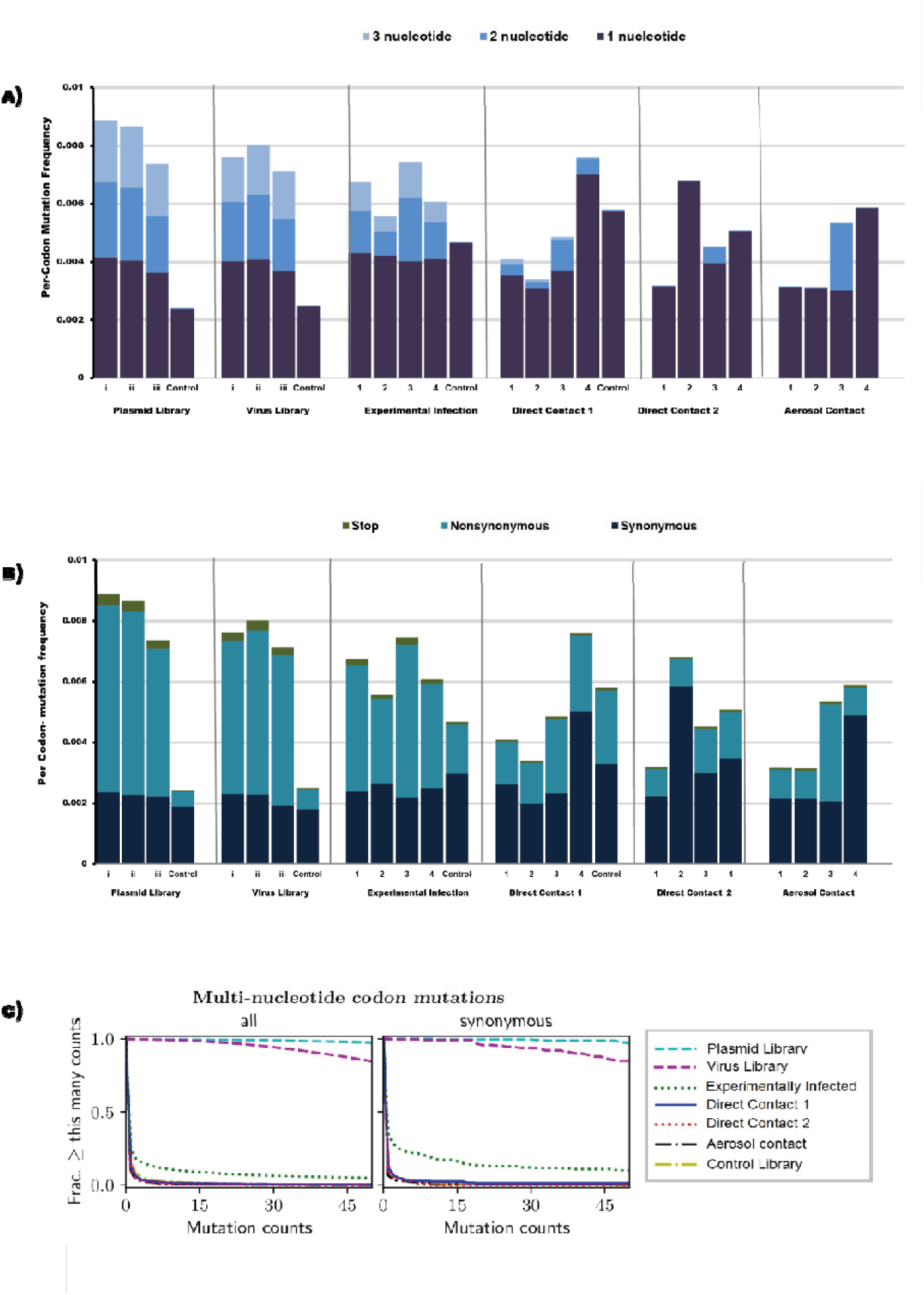
Following deep sequencing of the plasmid and virus libraries (and their controls), as well as from ferret nasal wash samples, the per-codon mutation frequency, composition and fraction of total mutations sampled in the viral NA were determined. **a)** The libraries were made of multi-nucleotide (2-or 3-) codon mutations, while the controls were not. Most viruses in direct contact 2 and aerosol contact animals contained single-nucleotide codon mutations. **b)** Viruses from ferret nasal wash samples generally had a greater ratio of synonymous changes to non-synonymous changes, indicating purifying selection. **c)** The fraction of multi-nucleotide mutations that were observed multiple times in the samples, after combining biological replicates, was >90% in the plasmid and virus libraries but was substantially reduced in ferret nasal wash samples.

In order to assess the completeness of the plasmid and virus libraries, the fraction of all multi-nucleotide codon mutations that were sampled multiple times was quantified (Figure 3C). Only multi-nucleotide mutations were considered as they are most likely to be introduced due to codon mutagenesis. In previous studies it was shown that to adequately sample 97% of all possible amino acids in a virus library, only 85% of all possible codon mutations needed to be present at least five times [62]. In our study, more than 99.5% of all multi-nucleotide codon mutations were sampled at least five times in the combined plasmid libraries and 99.2% were sampled at least five times in each individual replicate. In comparison, only 1.6% of all multi-nucleotide mutations were sampled five times in the control plasmid library. Similarly, the combined virus library had more than 99.0% of all multi nucleotide mutations sampled at least five times, with each individual replicate sampling at least 97% of all such mutations. The control virus sampled only 3.7% of such mutations at least five times. These results therefore indicate a high level of representation of all codon mutations in both the plasmid and virus libraries.

It should be noted that despite the large diversity of the NA genes in the plasmid and virus libraries, the frequency of each mutated codon in the library was low (0.0078-0.0087%) and the template NA sequence was overrepresented in codon counts.

### 3.3 Deep sequence analysis of ferret nasal wash samples reveal a stringent bottleneck at each transmission event restricting viral diversity

After confirming the completeness of the virus library in sampling all codon mutations, we aimed to investigate where replication and transmission in ferrets selected for fitter H275Y variants. This experiment was done in replicates of four (Figure 1). All animals in transmission lines 1, 2, 3, and 4, and control animals, were successfully infected as determined by shedding of detectable levels of virus in nasal wash samples (Figure 4A). In general, recipient or contact animals were found to shed detectable levels of virus within 24 hours post-exposure to their respective donors. A single nasal wash sample from each ferret was deep sequenced from one time point only (denoted by black arrows in Figure 4A).

**Figure 4:**
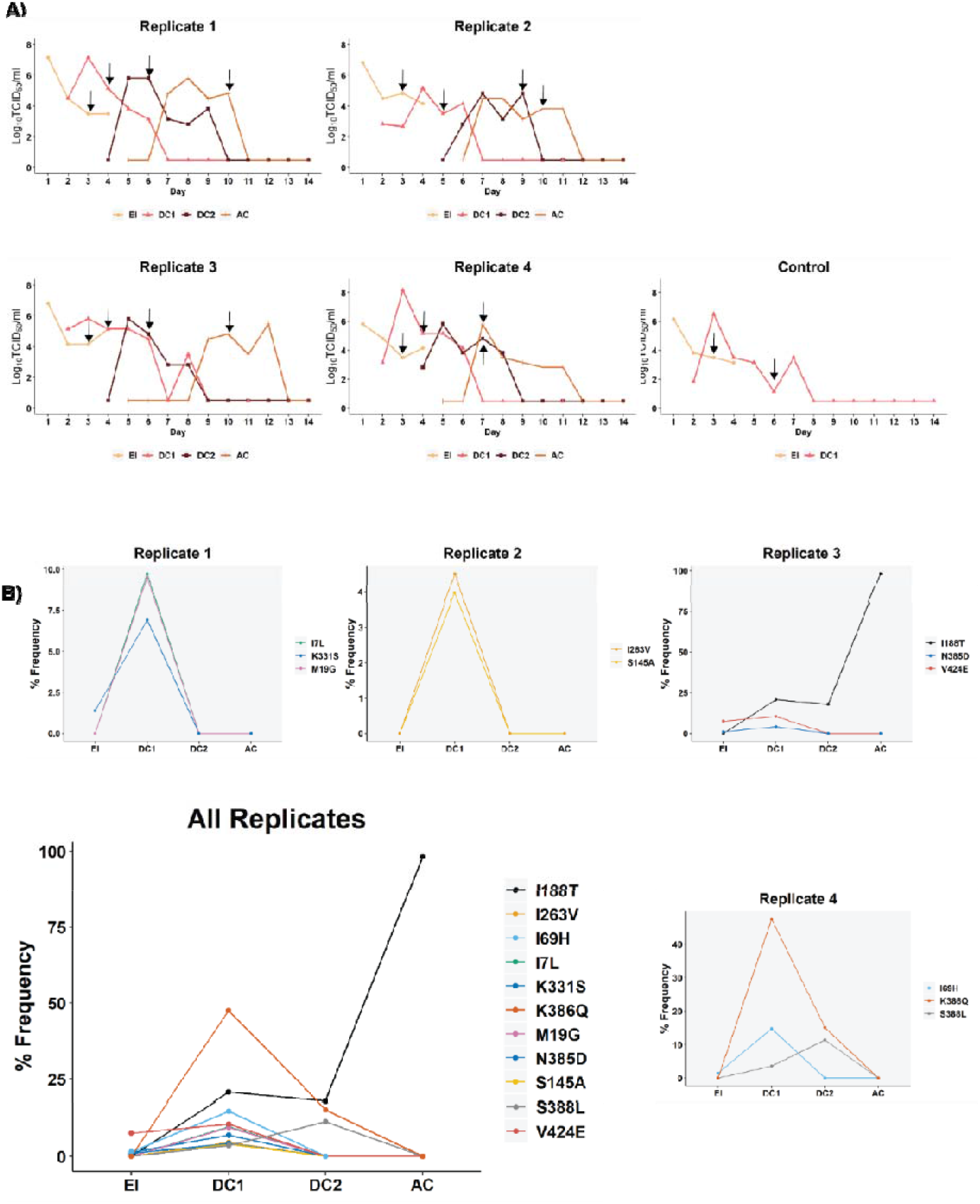
Viral titres and variant frequencies in nasal wash samples from ferrets experimentally infected with the NA-H275Y virus library, and from ferrets subsequently infected via transmission. **A)** At 24 hours post inoculation, experimentally infected animals were co-housed with direct contact 1 ferret. Nasal wash samples from direct contact 1 ferrets were monitored for infection, and, on the day that influenza infection was confirmed, they were cohoused with direct contact 2 ferrets. Nasal wash samples from direct contact 2 ferrets were monitored for infection, and, on the day that influenza infection was confirmed they were placed in a cage adjacent to aerosol contacts. All animals were nasal washed daily during the experiment and infectious virus was detected in nasal wash samples from animals along the transmission chain. For each animal, a single time-point (black arrows) was picked for analysis by deep sequencing. **B)** NGS data was aligned using Bowtie2 and variants observed at a greater than 1% frequency were called using VarScan, where the average read depth at each site was >7,000 and *p*-values for variant calls above 1% were <0.05 for all called positions. A different set of variants were observed in each transmission chain, and most variants were not observed beyond direct contact 1 animals. However, substitutions I188T, K386Q and S388L were present in direct contact 2 animals and are therefore of greater interest for further analysis.

The sequencing results from ferret nasal wash samples were aligned and analysed in two different ways: either using a combination of the mapmuts and dms_tools2 pipeline or aligned using the Bowtie2 program and screened for variants using Varscan. At least 10^6^ overlapping paired-end reads could be aligned to the NA genes using mapmuts and the read depth at each site was greater than 1.5 × 10^5^ reads per site. In contrast, at least 2.6 × 10^7^ could be aligned to the NA gene using Bowtie2 and >10^4^ reads per site were used to calculate mutation frequencies and *p*-values with Varscan.

There was a trend towards a reduced per-codon mutation frequency along the transmission chain (Figure 3A and B). Only 13.5%, 2.5%, 1.8% and 1.9% of all multi-nucleotide mutations were sampled in viruses from experimentally infected, direct contact 1, direct contact 2 and aerosol contact animals, respectively (Figure 3C). There was also a greater proportion of single-nucleotide, synonymous mutations and a reduced number of stop codons observed in the latter samples (Figure 3A and B). The composition of codon mutations in the animal infected with control virus consisted entirely of single-nucleotide substitutions (Figure 3A and B).

Nucleotide diversity in the viral populations was also analysed by calculating π from the Bowtie2 alignment data, which quantified the average number of pairwise differences per nucleotide site. The average π value of viruses from the experimentally infected animals (0.0016 ± 0.0003) was significantly higher than the average π values of viruses from direct contact 2 (π = 0.0006 ± 0.0004) and aerosol contact animals (π = 0.0004 ± 0.0003) (Table S1). Of note, no SNPs could be detected by VarScan in animals infected with only the control virus, SA16-H275Y, and therefore no π value is available for these animals.

The ratio between synonymous and nonsynonymous diversity, calculated by πN/πS, was also measured. In general πN/πS <1 indicates purifying selection that is purging deleterious mutations, πN/πS >1 indicates diversifying selection which favours new mutations and πN/πS =1 indicates neutrality [81]. With one exception, the ratio of πN/πS remained below 1 in viruses from all ferret nasal wash samples (Table S1).

Together, these results demonstrate that there is a significant reduction in viral diversity upon transmission of influenza virus in ferrets consistent with the presence of narrow bottleneck sizes during transmission. There is also evidence of purifying selection purging deleterious non-synonymous and stop mutations during virus replication in the ferrets. Of note, the H275Y substitution was not lost during transmission and remained fixed even in viruses from aerosol contact ferrets.

### 3.4 Bottleneck size estimate reveals a more stringent bottleneck during aerosol transmission than contact transmission

Given the results described above, it was of interest to learn more about the size of transmission bottlenecks (i.e. the number of transmitting viruses), as it was severely restricting viral diversity in recipient animals in our study. Utilising a mathematical model it was calculated that the approximate bottleneck sizes during contact transmission was somewhat varied between each transmission pairs with 23.87 viral particles being transmitted on average between ferrets (lower bound = 15, upper bound = 38) (Table 1). However, there was greater variability in estimates of bottleneck sizes during aerosol transmission, where an estimated 146 viral particles were transmitted between one pair (Replicate 1), while an average of 7.3 virus particles (lower bound = 3.7, upper bound = 13.7) were transmitted between the three other pairs of ferrets. With a more conservative minimum variant calling cut-off of 3%, the average number of particles being transmitted during contact exposure was 7.6 virus particles (lower bound= 2.8 and upper bound= 24.2) and for aerosol transmission was only 2 virus particles (lower bound=0.5 and upper bound=23, upper bound slightly skewed due to replicate 1).

**Table 1:** Bottleneck size estimated in donor: recipient pairs using the beta-binomial sampling method.

| Transmission route | Replicate | Donor<sup>a</sup> | Recipient<sup>b</sup> | Bottleneck Size (1% cut-off)<sup>c</sup> | Bottleneck Size (3% cut-off)<sup>d</sup> |
| --- | --- | --- | --- | --- | --- |
| Contact | Replicate 1 | EI | DC1 | 27 (18, 41) | 15 (7, 32) |
|  |  | DC1 | DC2 | 22 (11, 37) | 1 (0, 4) |
|  | Replicate 2 | EI | DC1 | 60 (38, 90) | 19 (5, 107) |
|  |  | DC1 | DC2 | 4 (2, 7) | 2 (1, 2) |
|  | Replicate 3 | EI | DC1 | 28 (18, 45) | 6 (3, 12) |
|  |  | DC1 | DC2 | 28 (17, 46) | 6 (2, 12) |
|  | Replicate 4 | EI | DC1 | 9 (6, 13) | 5 (2, 8) |
|  |  | DC1 | DC2 | 13 (6, 24) | 7 (3, 16) |
| Aerosol | Replicate 1 | DC2 | AC | 146 (73, 201) | 1 (0, 76) |
|  | Replicate 2 | DC2 | AC | 4 (2, 8) | 1 (0, 2) |
|  | Replicate 3 | DC2 | AC | 5 (2, 8) | 2 (1, 4) |
|  | Replicate 4 | DC2 | AC | 13 (7, 23) | 4 (1, 10) |

<sup>a</sup> EI- Experimentally infected, DC1- Direct Contact 1, DC2- Direct Contact 2
<sup>b</sup> AC- Aerosol contact
<sup>c</sup> Estimated size of bottleneck with lower and upper bounds when a minimum variant calling threshold is set at 1%
<sup>d</sup> Estimated size of bottleneck with lower and upper bounds when a minimum variant calling threshold is set at 3%

### 3.5 Amino acid substitutions under positive selection pressure in the presence of H275Y

Frequencies of nonsynonymous codon mutations (amino acid substitutions) across the transmission chain were analysed to see which variants increased in frequency following transmission (Figure 4B). Variants that increased in frequency following transmission are likely to be under positive selection pressure and hence contain substitutions that may be permissive for H275Y. It should be noted here that each variant in the original library was present at very low levels (0.008%), and competing with several thousand other variants, and therefore even modest increases in frequency to 4-5% can be indicative of a positive selection pressure.

The results reveal that each replicate of the transmission chain was different from the other (Figure 4B). In replicate 1, M19G (ATG> GGC) and I7L (ATA >CTC) were observed at frequencies of 7-9% in direct contact 1 ferrets, despite being below the 1% detection threshold in experimentally infected animals. The substitution K331S (AAG>TCG) was observed at 1.4% in experimentally infected animals and rose to a frequency of 6.9% in direct contact 1 animals. Interestingly, the K331S substitution (AAG>TCG) was also observed at frequencies of 1.4-1.6% experimentally infected animals from replicates 3 and 4, but these viruses did not transmit to their corresponding recipients.

In replicate 2, substitutions I263V (ATA>GTA) and S145A (TCC>GCG) were observed at 3-4% in direct contact 1 ferrets, but were both lost subsequently down the transmission chain. Similarly, in replicate 3 the V424E (GTT>GAG) substitution increased from 7.6% in experimentally infected animals to 10% in direct contact 1 animals but was not observed in nasal wash samples later in the transmission chain. The V424E substitution was also observed at a frequency of 5.8% in replicate 4 animals after experimental infection but did not transmit to the corresponding contact animal. The substitution N385D (AAT>GAC) In replicate 3, increased from a frequency of 1% in experimentally infected animal to 4.21% in direct contact 1 animals before disappearing altogether. In contrast, the I188T (ATC>ACG) increased from less than 1% in experimentally infected animals to 17-20% in direct contact 1 and 2 animals and to 98% in aerosol contact animals.

In replicate 4, substitution K386Q (AAA>CAA) increased to 47.5% in direct contact 1 animals but was reduced to 15% in direct contact 2 animals, and then lost altogether in aerosol contact animals. The substitution S388L (TCA>TTA) was observed at 3.5% in direct contact 1 animals and 11% in direct contact 2 animals but not in aerosol contact animals. Finally, the I69H (ATC>CAC) substitution increased from 1.5% in experimentally infected animals to 14.7% in direct contact 1 animals, before being lost in subsequent animals along the transmission chain. Of note, the I69H substitution was observed in all experimentally infected animals (3% in replicate 1, 1.6% in replicate 2 and 4.5% in replicate 3) but only transmitted to direct contact 1 animals in replicate 4.

As most substitutions were lost following transmission from direct contact 1 animals, it was of interest to sequence nasal washes of direct contact 1 animal across different experimental days to test for the genetic stability of the variants observed in these animals (Supplementary Figure S2). The results showed all the variants observed were stable during the experimental days in direct contact 1 animals, and further that V424E increased in frequency from 4% to 19% in replicate 3 direct contact 1 animals. This analysis also revealed two more substitutions in replicate 1 direct contact 1 animal, D451G (GAC>CTG) and Y402A (TAT>GCG), present at 37% and 25% respectively, that were below detection limit in its corresponding experimentally infected animal.

Full genome sequencing revealed that the reversion of the S31N mutation remained stable during transmission events, and while a small number of variants were observed, no sustained changes in the internal genes of the virus was seen (Table S2).

### 3.6 Evaluation of SA16-H275Y fitness with I188T, K386Q and S388L

As substitutions I188T, K386Q and S388L were present in direct contact 2 animals, they were analysed further for their effect on enzyme function in the presence of the H275Y NA substitution. The I188T substitution was of particular interest as it reached a frequency of approximately 98% in the replicate 3 aerosol contact animal. Significant variability was observed in the NA activity assay with a relative NA activity of 77 ± 21% recorded for the H275Y-NA (Figure 6A). The impact of all candidate substitutions on NA activity and expression was compared to that of the H275Y-NA. Overall, there was a trend towards increased activity in H275Y+I188T-NA and H275Y+S388L NA, with relative NA activities of 116 ± 84% and 87 ± 54% respectively; however, these increases were not significant. The H275Y+K386Q-NA showed similar levels of relative NA activity (79 ± 38%) to the H275Y-NA.

**Figure 6:**
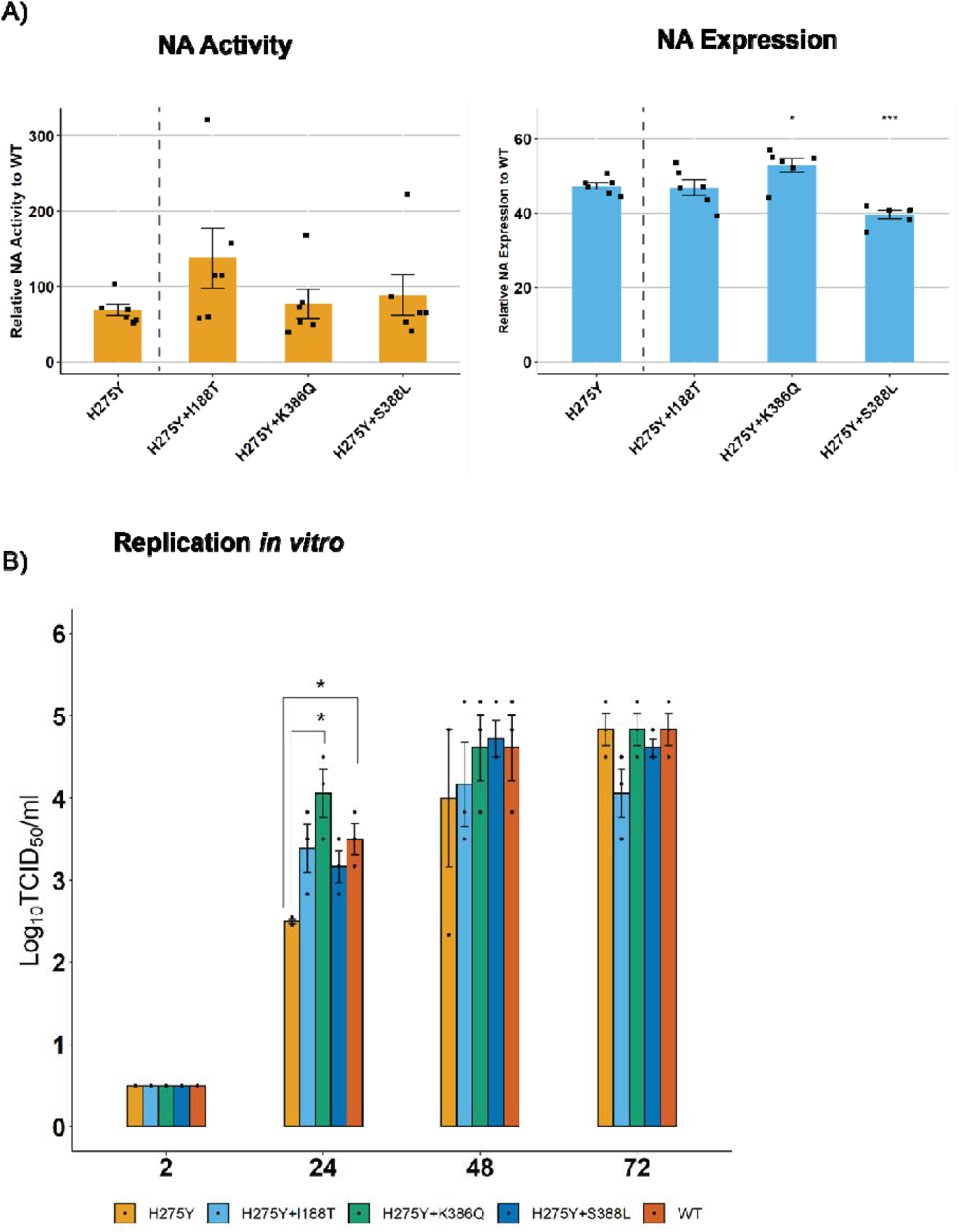
Relative NA activity and expression of variant NA glycoproteins with candidate substitutions identified from the experimental approaches described were determined and compared to the H275Y-NA. The NA glycoprotein of the A/South Australia/16/2017 virus was mutated such that it contained the H275Y substitution by itself or in combinations with candidate permissive substitutions. The proteins were expressed in cells following transfection of 293T cells and the relative NA activity and expression were calculated as a percentage of wild-type (WT) NA protein (lacking any substitution). The assay was performed in duplicate on three independent occasions and the mean ± SD are shown. The relative NA activity and expression for the NA proteins containing candidate substitutions was compared against that of the H275Y-NA using a Student ‘s unpaired two-tailed *t*-test. * p<0.05, ** p<0.01 **b)** The replication kinetics of SA16-H275Y, SA16-WT and SA16-H275Y modified with either I188T, K386Q or S388L NA substitution was measured in A549 cells infected with an MOI 0.1. The experiment was performed in triplicates and viral titres at each time point were measured using a Student ‘s unpaired two-tailed *t*-test. * p<0.05, ** p<0.01.

The relative NA expression of the H275Y+I188T-NA was similar to that of the H275Y-NA (46 ± 4 % vs 44 ± 5%). However, relative NA expression was significantly increased in the H275Y+K386Q-NA (50 ± 5%) compared to the H275Y-NA. Conversely, relative NA expression was significantly reduced with the H275Y+S388L (40 ± 2%) compared to the H275Y-NA (Figure 6A).

The substitutions I188T, K386Q and S388L were studied further in an *in vitro* replication kinetics experiment (Figure 6B). All three substitutions led to moderate improvements in viral titres compared to the SA16-H275Y virus at 24 and 48 hrs post infection, with a significant increase in virus titres observed with the SA16-H275Y+K386Q at 24 hr post infection compared to the SA16-H275Y virus (4.0 ± 0.3 Log_10_ TCID_50_/ml vs 2.5 ± 0.02 Log_10_ TCID_50_/ml).

## 4. Discussion

This study explored two different approaches to predict neuraminidase substitutions that may be potentially permissive for the H275Y NA substitution, which is known to reduce the susceptibility of A(H1N1)pdm09 viruses to oseltamivir. The first approach utilised computational analyses to predict *in silico* protein stability (based on free energy change) and proposed candidate substitutions S95N, S286G and S299A as potentially permissive for H275Y. Analysis of all NA sequences in the GISAID database (34,510 sequences) (Figure S3), show that these substitutions occur at a low frequency in natural sequences. *In vitro* experimental analysis was not able to confirm that these substitutions substantially improved relative NA activity, although the reduction in NA expression due to H275Y was offset by 10% with a combination of S95N+S286G+S299A substitutions. However, it was also observed that the combination of S95N+S286G+S299A offset loss in virus titres due to H275Y during *in vitro* replication, suggesting they may play a permissive role if studied in more depth in future studies, utilising *in vivo* models.

The second experimental approach utilised a virus library representing all single NA amino acid substitutions (except H275Y, which was fixed) to select for fit variants during serial transmission in ferrets. A somewhat similar strategy was previously utilised by Wu *et al*., whereby error-prone PCR was used to generate a virus library with the H275Y substitution, and fit variants were selected after cell-culture passaging [82]. However, a number of important differences distinguish our study from that of Wu *et al*. First, we have used a contemporary virus strain (A/South Australia/16/2017 vs A/WSN/33 [82]) and performed mutagenesis at a codon level instead at a single nucleotide level. Moreover, we utilised an animal model to select for variants with high transmission and replication fitness, instead of cell culture passaging to select for variants with high replicative fitness [82].

In our study, strong purifying selection was observed in the experimentally infected animals and the stringent transmission bottleneck severely restricted the viral diversity in the recipient animals. The transmission bottleneck for contact transmission was estimated to allow between 14-37 virus particles to transmit between ferrets using a less conservative sequence analysis threshold, and between 2-24 virus particles with a more conservative sequence analysis threshold. The bottleneck was more stringent during aerosol transmission (conservative estimate: 1-5 virus particles, less conservative estimate: 4-13 virus particles), though there was an outlier in replicate 1. Our estimates were similar to those proposed by previous experiments in ferret and guinea pig models [83, 84] and in a human household transmission study [85]. However, a previous analysis of datasets from human household transmission studies have proposed a much looser transmission bottleneck (146-200 virus particles) [77]

The stochastic nature of transmission events was a limitation of the experimental approach, as a different set of variants were seen to pass from the experimentally infected animals to their direct contact recipients in each of the 4 transmission chains. Amongst the variants observed in these replicates, substitutions I7L and M19G occurred in the transmembrane region of the NA protein, while I69H occurred in the linker region connecting the transmembrane region to the catalytic domain [86]. The remaining substitutions are in the catalytic head of the NA protein. None of these variants were selected by cell culture passaging in the previous study by Wu *et al*. [82].

The substitutions I188T, K386Q and S388L were of greater interest as they were detected in nasal wash samples after two transmission events. However, characterising the effect of these substitutions in an NA enzyme function showed that K386Q offset the loss in NA expression due to H275Y by only 5% and, while there was a trend for improved NA activity with I188T and S388L it was not significant. Modest improvements in virus titres were however observed in reverse genetics viruses with all three substitutions in combination with H275Y, compared to viruses with the H275Y substitution alone.

Interestingly, sequence database analysis of the influenza viral NA revealed that the substitution I188T has increased in frequency from 1.1% in circulating viruses in 2016 to 98% in 2020 (Figure S3). The strong selection for this substitution in at least one of our replicates, in an early (January) 2017 virus, suggests a degree of predictive capability in our experimental analysis. While the high prevalence of I188T in currently circulating viruses suggests that this substitution is unlikely to be fully permissive for H275Y, since H275Y prevalence has not increased since 2016, this substitution is still of interest for further study in combination with other candidate substitutions.

The K386Q substitution is also of great interest, because even though it has been observed only once in natural influenza sequences, substitutions at amino acid position 386 are common and have been suggested as candidates for permissive substitutions in previous studies. For example, a N386S substitution was observed in the NA of viruses from the cluster of H275Y A(H1N1)pdm09 variants in Hunter New England in 2011 although it was not present in the majority of the strains circulating worldwide that year [52, 54]. The N386S substitution did not improve the loss in NA expression or activity due to H275Y and was not studied further in ferrets [54]. In a predictive study by Bloom *et al*. computational analyses suggested that N386E may be a potentially permissive substitution, however this substitution was not found to improve NA activity or expression [56]. Interestingly, the computational analysis in this study also predicated substitutions at position 386, namely N386S and N386D, to improve *in silico* protein stability (Figure S1). Finally, the N386K substitution was observed in a cluster of H275Y variants in Sapporo, Hokkaido, Japan in 2014, and the lysine (K) has been since incorporated in all circulating strains [53]. The K386Q substitution therefore needs to be verified further, to assess its effects on virus fitness, in future *in vivo* studies.

Amongst the other candidate substitutions proposed by the experimental approach, I7L, M19G, I69H, S145A and V424E were not observed in the natural influenza sequences. However, the frequency of the I263V substitution has increased from 0.4% in 2017 to 3.4% in 2019 (Figure S4) and the S388L substitution is observed at very low frequencies in most years.

The approaches developed in this study provide opportunities for a number of lines of additional work. For example, we did not analyse synonymous substitutions in our studies although we did note that certain synonymous mutations increased in frequency following transmission in ferrets, suggesting that some of these may have had a beneficial effect on viral fitness. Previous experiments studying the influenza A virus HA glycoprotein have demonstrated that synonymous mutations can have an impact on experimental viral fitness [87]. There is also the opportunity to combine the inferences from our experimental approach with our computational approach to narrow down on permissive substitutions. Finally, we can explore alternate approaches to select for fit variants, such as passaging in representative human cell lines like the differentiated normal human bronchial epithelial cells or airway organoids [88, 89], or by passaging a wild-type virus library (instead of a library where H275Y is fixed), in increasing oseltamivir pressure.

In summary, after developing two different approaches, this study proposed a number of candidate substitutions that may be potentially permissive for H275Y. A selection of these substitutions was tested for their ability to compensate for loss of NA enzyme function, however only moderate improvements were observed. A smaller subset of these selected substitutions was studied for their impact on *in vitro* virus replication, and all were found to improve replication fitness of the H275Y containing virus up to a certain degree. It remains important to analyse the possible impact of these substitution in an *in vivo* model in future studies. This is especially true for the substitutions that have been observed to increase in frequency in the influenza database in recent years, such as I188T, which may be one step forward for the virus to become fully permissive for H275Y. Together, the different approaches utilised in this paper provides insights into the fitness landscape of H275Y variant influenza A(H1N1) viruses and presents opportunities for further work, including tools for future experiments aimed at understanding virus evolution in-depth.

## Acknowledgements

We thank Rebecca Bowyer, Thomas Cumming, Anthony Gomes, Simone Azzopardi, Charlene Plasencia and Danijela Krmek at Bio-resource facility of University of Melbourne for providing assistance in animal handling. We also thank Dr. Michelle Wille in providing help in setting up bioinformatics analysis through the server and Mr. Leo Lee for providing assistance during animal experiments. Finally, this project would not have been possible without the generous support and guidance of Dr. Jesse Bloom whose protocols and pipelines were utilised heavily throughout this manuscript for creating and analysing virus libraries.

The Melbourne WHO Collaborating Centre for Reference and Research on Influenza is supported by the Australian Government Department of Health.

## References

1. Gubareva, L.V., et al., Global update on the susceptibility of human influenza viruses to neuraminidase inhibitors, 2015–2016. Antiviral research, 2017. 146: p. 12–20.

2. Hurt, A.C., et al., Global update on the susceptibility of human influenza viruses to neuraminidase inhibitors, 2014-2015. Antiviral Res, 2016. 132: p. 178–85.

3. Takashita, E., et al., Global update on the susceptibility of human influenza viruses to neuraminidase inhibitors, 2013-2014. Antiviral Res, 2015. 117: p. 27–38.

4. Meijer, A., et al., Global update on the susceptibility of human influenza viruses to neuraminidase inhibitors, 2012-2013. Antiviral Res, 2014. 110: p. 31–41.

5. Lackenby, A., et al., Global update on the susceptibility of human influenza viruses to neuraminidase inhibitors and status of novel antivirals, 2016-2017. Antiviral Res, 2018. 157: p. 38–46.

6. von Itzstein, M., The war against influenza: discovery and development of sialidase inhibitors. Nat Rev Drug Discov, 2007. 6(12): p. 967–74.

7. Itzstein, M.v., et al., Rational design of potent sialidase-based inhibitors of influenza virus replication. Nature, 1993(6428): p. 418.

8. Ferraris, O. and B. Lina, Mutations of neuraminidase implicated in neuraminidase inhibitors resistance. J Clin Virol, 2008. 41(1): p. 13–9.

9. Gubareva, L.V., Molecular mechanisms of influenza virus resistance to neuraminidase inhibitors. Virus Res, 2004. 103(1-2): p. 199–203.

10. Tashiro, M., et al., Surveillance for neuraminidase-inhibitor-resistant influenza viruses in Japan, 1996-2007. Antivir Ther, 2009. 14(6): p. 751–61.

11. Sheu, T.G., et al., Surveillance for neuraminidase inhibitor resistance among human influenza A and B viruses circulating worldwide from 2004 to 2008. Antimicrob Agents Chemother, 2008. 52(9): p. 3284–92.

12. Wang, M.Z., C.Y. Tai, and D.B. Mendel, Mechanism by which mutations at his274 alter sensitivity of influenza a virus n1 neuraminidase to oseltamivir carboxylate and zanamivir. Antimicrob Agents Chemother, 2002. 46(12): p. 3809–16.

13. Collins, P.J., et al., Structural basis for oseltamivir resistance of influenza viruses. Vaccine, 2009. 27(45): p. 6317–23.

14. Collins, P.J., et al., Crystal structures of oseltamivir-resistant influenza virus neuraminidase mutants. Nature, 2008. 453(7199): p. 1258–61.

15. Kawai, N., et al., Clinical effectiveness of oseltamivir and zanamivir for treatment of influenza A virus subtype H1N1 with the H274Y mutation: a Japanese, multicenter study of the 2007-2008 and 2008-2009 influenza seasons. Clin Infect Dis, 2009. 49(12): p. 1828–35.

16. Mungall, B.A., X. Xu, and A. Klimov, Surveillance of influenza isolates for susceptibility to neuraminidase inhibitors during the 2000-2002 influenza seasons. Virus Res, 2004. 103(1-2): p. 195–7.

17. Monto, A.S., et al., Detection of influenza viruses resistant to neuraminidase inhibitors in global surveillance during the first 3 years of their use. Antimicrob Agents Chemother, 2006. 50(7): p. 2395–402.

18. Escuret, V., et al., Detection of human influenza A (H1N1) and B strains with reduced sensitivity to neuraminidase inhibitors. J Clin Virol, 2008. 41(1): p. 25–8.

19. Monitoring of neuraminidase inhibitor resistance among clinical influenza virus isolates in Japan during the 2003-2006 influenza seasons. Wkly Epidemiol Rec, 2007. 82(17): p. 149–50.

20. Carr, J., et al. Virological assessment in vitro and in vivo of an influenza H1N1 virus with a H274Y mutation in the neuraminidase gene. in Antiviral research. 2000. LSEVIER SCIENCE BV PO BOX 211, 1000 AE AMSTERDAM, NETHERLANDS.

21. Ives, J.A.L., et al., The H274Y mutation in the influenza A/H1N1 neuraminidase active site following oseltamivir phosphate treatment leave virus severely compromised both in vitro and in vivo. Antiviral research, 2002. 55(2): p. 307–317.

22. Herlocher, M.L., et al., Influenza viruses resistant to the antiviral drug oseltamivir: transmission studies in ferrets. J Infect Dis, 2004. 190(9): p. 1627–30.

23. Abed, Y., N. Goyette, and G. Boivin, A reverse genetics study of resistance to neuraminidase inhibitors in an influenza A/H1N1 virus. Antivir Ther, 2004. 9(4): p. 577–81.

24. Baz, M., Y. Abed, and G. Boivin, Characterization of drug-resistant recombinant influenza A/H1N1 viruses selected in vitro with peramivir and zanamivir. Antiviral Res, 2007. 74(2): p. 159–62.

25. Lackenby, A., et al., Emergence of resistance to oseltamivir among influenza A(H1N1). Peer-reviewed European information on communicable disease surveillance and control, 2008: p. 113.

26. Meijer, A., et al., Oseltamivir-resistant influenza virus A (H1N1), Europe, 2007–08 season. 2009.

27. Hauge, S.H., et al., Oseltamivir-Resistant Influenza Viruses A (H1N1), Norway, 2007–08. Emerging Infectious Diseases, 2009. 15(2): p. 155–162.

28. Moscona, A., Global Transmission of Oseltamivir-Resistant Influenza. New England Journal of Medicine, 2009. 360(10): p. 953–956.

29. Besselaar, T.G., et al., Widespread Oseltamivir Resistance in Influenza A Viruses (H1N1), South Africa. Emerging Infectious Diseases, 2008. 14(11): p. 1809–1810.

30. Dawood, F.S., et al., Emergence of a novel swine-origin influenza A (H1N1) virus in humans. N Engl J Med, 2009. 360(25): p. 2605–15.

31. Michaelis, M., H.W. Doerr, and J. Cinatl, Jr., An influenza A H1N1 virus revival -pandemic H1N1/09 virus. Infection, 2009. 37(5): p. 381–9.

32. Gubareva, L.V., et al., Comprehensive assessment of 2009 pandemic influenza A (H1N1) virus drug susceptibility in vitro. Antivir Ther, 2010. 15(8): p. 1151–9.

33. Baz, M., et al., Effect of the neuraminidase mutation H274Y conferring resistance to oseltamivir on the replicative capacity and virulence of old and recent human influenza A(H1N1) viruses. J Infect Dis, 2010. 201(5): p. 740–5.

34. Bouvier, N.M., S. Rahmat, and N. Pica, Enhanced mammalian transmissibility of seasonal influenza A/H1N1 viruses encoding an oseltamivir-resistant neuraminidase. J Virol, 2012. 86(13): p. 7268–79.

35. Hurt, A.C., et al., Assessing the Viral Fitness of Oseltamivir-Resistant Influenza Viruses in Ferrets, Using a Competitive-Mixtures Model. Journal of Virology, 2010. 84(18): p. 9427–9438.

36. Duan, S., et al., Epistatic interactions between neuraminidase mutations facilitated the emergence of the oseltamivir-resistant H1N1 influenza viruses. Nat Commun, 2014. 5.

37. Bloom, J.D., L.I. Gong, and D. Baltimore, Permissive Secondary Mutations Enable the Evolution of Influenza Oseltamivir Resistance. Science, 2010. 328(5983): p. 1272–1275.

38. Abed, Y., et al., Role of permissive neuraminidase mutations in influenza A/Brisbane/59/2007-like (H1N1) viruses. PLoS Pathog, 2011. 7(12): p. 8.

39. Abed, Y., et al., Permissive changes in the neuraminidase play a dominant role in improving the viral fitness of oseltamivir-resistant seasonal influenza A(H1N1) strains. Antiviral Res, 2015. 114: p. 57–61.

40. Rameix-Welti, M.A., et al., Neuraminidase of 2007-2008 influenza A(H1N1) viruses shows increased affinity for sialic acids due to the D344N substitution. Antivir Ther, 2011. 16(4): p. 597–603.

41. Rameix-Welti, M.A., et al., Enzymatic properties of the neuraminidase of seasonal H1N1 influenza viruses provide insights for the emergence of natural resistance to oseltamivir. PLoS Pathog, 2008. 4(7): p. 1000103.

42. Ginting, T.E., et al., Amino Acid Changes in Hemagglutinin Contribute to the Replication of Oseltamivir-Resistant H1N1 Influenza Viruses. Journal of Virology, 2012. 86(1): p. 121–127.

43. Takashita, E., et al., Global update on the susceptibilities of human influenza viruses to neuraminidase inhibitors and the cap-dependent endonuclease inhibitor baloxavir, 2017-2018. Antiviral Res, 2020. 175: p. 104718.

44. Hamelin, M.-È., et al., Oseltamivir-Resistant Pandemic A/H1N1 Virus Is as Virulent as Its Wild-Type Counterpart in Mice and Ferrets. PLoS Pathogens, 2010. 6(7): p. e1001015.

45. Wong, D.D.Y., et al., Comparable Fitness and Transmissibility between Oseltamivir-Resistant Pandemic 2009 and Seasonal H1N1 Influenza Viruses with the H275Y Neuraminidase Mutation. Journal of Virology, 2012. 86(19): p. 10558–10570.

46. Kiso, M., et al., Characterization of oseltamivir-resistant 2009 H1N1 pandemic influenza A viruses. PLoS Pathog, 2010. 6(8): p. 1001079.

47. Seibert, C.W., et al., Oseltamivir-Resistant Variants of the 2009 Pandemic H1N1 Influenza A Virus Are Not Attenuated in the Guinea Pig and Ferret Transmission Models. Journal of Virology, 2010. 84(21): p. 11219–11226.

48. Memoli, M.J., et al., Multidrug-resistant 2009 pandemic influenza A(H1N1) viruses maintain fitness and transmissibility in ferrets. J Infect Dis, 2011. 203(3): p. 348–57.

49. Duan, S., et al., Oseltamivir–Resistant Pandemic H1N1/2009 Influenza Virus Possesses Lower Transmissibility and Fitness in Ferrets. PLoS Pathogens, 2010. 6(7): p. e1001022.

50. Brookes, D.W., et al., Pandemic H1N1 2009 influenza virus with the H275Y oseltamivir resistance neuraminidase mutation shows a small compromise in enzyme activity and viral fitness. J Antimicrob Chemother, 2011. 66(3): p. 466–70.

51. Pinilla, L.T., et al., The H275Y neuraminidase mutation of the pandemic A/H1N1 influenza virus lengthens the eclipse phase and reduces viral output of infected cells, potentially compromising fitness in ferrets. J Virol, 2012. 86(19): p. 10651–60.

52. Hurt, A.C., et al., Characteristics of a widespread community cluster of H275Y oseltamivir-resistant A(H1N1)pdm09 influenza in Australia. J Infect Dis, 2012. 206(2): p. 148–57.

53. Takashita, E., et al., Characterization of a large cluster of influenza A(H1N1)pdm09 viruses cross-resistant to oseltamivir and peramivir during the 2013-2014 influenza season in Japan. Antimicrob Agents Chemother, 2015. 59(5): p. 2607–17.

54. Butler, J., et al., Estimating the fitness advantage conferred by permissive neuraminidase mutations in recent oseltamivir-resistant A(H1N1)pdm09 influenza viruses. PLoS Pathog, 2014. 10(4).

55. Abed, Y., et al., Impact of potential permissive neuraminidase mutations on viral fitness of the H275Y oseltamivir-resistant influenza A(H1N1)pdm09 virus in vitro, in mice and in ferrets. J Virol, 2014. 88(3): p. 1652–8.

56. Bloom, J.D., J.S. Nayak, and D. Baltimore, A Computational-Experimental Approach Identifies Mutations That Enhance Surface Expression of an Oseltamivir-Resistant Influenza Neuraminidase. PLoS ONE, 2011. 6(7): p. e22201.

57. Abed, Y., et al., Comparison of early and recent influenza A(H1N1)pdm09 isolates harboring or not the H275Y neuraminidase mutation, in vitro and in animal models. Antiviral Res, 2018. 159: p. 26–34.

58. Schymkowitz, J., et al., The FoldX web server: an online force field. Nucleic Acids Res, 2005. 33(Web Server issue).

59. Eswar, N., et al., Comparative protein structure modeling using Modeller. Curr Protoc Bioinformatics, 2006. 5(5).

60. Thyagarajan, B. and J.D. Bloom, The inherent mutational tolerance and antigenic evolvability of influenza hemagglutinin. Elife, 2014. 8(3): p. 03300.

61. Bloom, J.D., An experimentally determined evolutionary model dramatically improves phylogenetic fit. Mol Biol Evol, 2014. 31(8): p. 1956–78.

62. Doud, M.B. and J.D. Bloom, Accurate Measurement of the Effects of All Amino-Acid Mutations on Influenza Hemagglutinin. Viruses, 2016. 8(6).

63. Hoffmann, E., et al., Universal primer set for the full-length amplification of all influenza A viruses. Arch Virol, 2001. 146(12): p. 2275–89.

64. Hoffmann, E., et al., A DNA transfection system for generation of influenza A virus from eight plasmids. Proc Natl Acad Sci U S A, 2000. 97(11): p. 6108–13.

65. Lee, J.M., et al., Deep mutational scanning of hemagglutinin helps predict evolutionary fates of human H3N2 influenza variants. Proc Natl Acad Sci U S A, 2018. 115(35): p. E8276–E8285.

66. Ramakrishnan, M.A., Determination of 50% endpoint titer using a simple formula. World journal of virology, 2016. 5(2): p. 85–86.

67. Pedersen, J.C., Hemagglutination-inhibition assay for influenza virus subtype identification and the detection and quantitation of serum antibodies to influenza virus. Methods Mol Biol, 2014: p. 0758-8_2.

68. Oh, D.Y. and A.C. Hurt, Using the ferret as an animal model for investigating influenza antiviral effectiveness. Frontiers in Microbiology, 2016. 7.

69. Oh, D.Y., et al., Evaluation of oseltamivir prophylaxis regimens for reducing influenza virus infection, transmission and disease severity in a ferret model of household contact. J Antimicrob Chemother, 2014. 69(9): p. 2458–69.

70. McCrone, J.T. and A.S. Lauring, Measurements of Intrahost Viral Diversity Are Extremely Sensitive to Systematic Errors in Variant Calling. J Virol, 2016. 90(15): p. 6884–95.

71. Zhou, B., et al., Single-reaction genomic amplification accelerates sequencing and vaccine production for classical and Swine origin human influenza a viruses. Journal of virology, 2009. 83(19): p. 10309–10313.

72. Katoh, K. and D.M. Standley, MAFFT multiple sequence alignment software version 7: improvements in performance and usability. Molecular biology and evolution, 2013. 30(4): p. 772–780.

73. Delano, W., L, PyMol, D. Scientific, Editor. 2002, DeLano Scientific, San Carlos, CA, 700.

74. van der Vries, E., et al., H1N1 2009 pandemic influenza virus: resistance of the I223R neuraminidase mutant explained by kinetic and structural analysis. PLoS Pathog, 2012. 8(9): p. 20.

75. Koboldt, D.C., et al., VarScan: variant detection in massively parallel sequencing of individual and pooled samples. Bioinformatics (Oxford, England), 2009. 25(17): p. 2283–2285.

76. Nelson, C.W., L.H. Moncla, and A.L. Hughes, SNPGenie: estimating evolutionary parameters to detect natural selection using pooled next-generation sequencing data. Bioinformatics (Oxford, England), 2015. 31(22): p. 3709–11.

77. Sobel Leonard, A., et al., Transmission Bottleneck Size Estimation from Pathogen Deep-Sequencing Data, with an Application to Human Influenza A Virus. Journal of Virology, 2017. 91(14): p. e00171–17.

78. Poon, L.L., et al., Quantifying influenza virus diversity and transmission in humans. Nat Genet, 2016. 48(2): p. 195–200.

79. Farrukee, R., et al., Characterization of substitutions in the neuraminidase of A (H7N9) influenza viruses selected following serial passage in the presence of different neuraminidase inhibitors. Antiviral research, 2019.

80. Farrukee, R., et al., Characterization of Influenza B Virus Variants with Reduced Neuraminidase Inhibitor Susceptibility. Antimicrob Agents Chemother, 2018. 62(11): p. 01081–18.

81. Nelson, C.W., L.H. Moncla, and A.L. Hughes, SNPGenie: estimating evolutionary parameters to detect natural selection using pooled next-generation sequencing data. Bioinformatics (Oxford, England), 2015. 31(22): p. 3709–3711.

82. Wu, N.C., et al., Systematic identification of H274Y compensatory mutations in influenza A virus neuraminidase by high-throughput screening. J Virol, 2013. 87(2): p. 1193–9.

83. Frise, R., et al., Contact transmission of influenza virus between ferrets imposes a looser bottleneck than respiratory droplet transmission allowing propagation of antiviral resistance. Sci Rep, 2016. 6(29793).

84. Varble, A., et al., Influenza A virus transmission bottlenecks are defined by infection route and recipient host. Cell Host Microbe, 2014. 16(5): p. 691–700.

85. McCrone, J.T., et al., Stochastic processes constrain the within and between host evolution of influenza virus. Elife, 2018. 7: p. e35962.

86. Maurer-Stroh, S., et al., Mapping the sequence mutations of the 2009 H1N1 influenza A virus neuraminidase relative to drug and antibody binding sites. Biology Direct, 2009. 4(1): p. 18.

87. Canale, A.S., et al., Synonymous Mutations at the Beginning of the Influenza A Virus Hemagglutinin Gene Impact Experimental Fitness. J Mol Biol, 2018. 430(8): p. 1098–1115.

88. Zhou, J., et al., Differentiated human airway organoids to assess infectivity of emerging influenza virus. Proc Natl Acad Sci U S A, 2018. 115(26): p. 6822–6827.

89. Davis, A.S., et al., Validation of normal human bronchial epithelial cells as a model for influenza A infections in human distal trachea. J Histochem Cytochem, 2015. 63(5): p. 312–28.

