## Supplementary Text S1 for "Predicting potentially permissive substitutions that improve the fitness of A(H1N1)pdm09 viruses bearing the H275Y NA substitution"

Supplementary Text S1: Details for making Virus Library

*Codon Mutagenesis PCR*

Codon-based mutagenesis was used to generate a NA plasmid library as has been previously described for influenza A virus HA and NP genes [60-62]. Briefly, mutagenesis primers were designed to introduce randomized NNN nucleotide triplets at each codon site (470 forward and reverse primers were ordered for the 470-residue NA gene), although primers overlapping the H275Y site were not included. The forward and reverse primers were mixed in equimolar ratios to make forward and reverse primer pools. Prior to mutagenesis, template NA from a A(H1N1)pdm09 virus (A/South Australia/16/2017) which expressed the H275Y substitution (H275Y-NA) was amplified from pHW2000-NA plasmid using the Platinum™ *Taq* DNA Polymerase High Fidelity kit (Thermo Fisher Scientific, Australia) and the 5’ For-N1 (AACCGGAGTACTGGTCGACCTCCG) and 3’ Rev-N1 (GGTATATCTTTCGCTCCGAGTCGGC) primers. PCR products were purified using the QIAquick PCR Purification Kit (Qiagen, Germany). Two low-cycle PCR reactions were carried out on the template NA, one with the forward pooled primers and another with the reverse pooled primers. The PCR reactions contained 15 µL 2x KOD Hot Start Master mix (Merck, USA), 2 µL of either forward or reverse primer pool at final concentration of 4.5 µM, 2 µL of either 5’ For-N1 or 3’ Rev-N1 primer at 4.5 µM, 4 µL template at 3 ng/µL and 7µL water.

The PCR cycling conditions were as follows:

1. 95 ͦ C for 2:00
2. 95 ͦ C for 0:20
3. 70 ͦ C for 0:01
4. 50 ͦ C for 0:30, cooling to 50 C at 0.5 C/second
5. 70 ͦ C for 0:40
6. Repeat steps 2) to 5) for 6 additional cycles

Following PCR, the forward and reverse fragments were diluted 1:4 in elution buffer and used in a joining reaction. The joining reaction contained 15 µL 2X KOD Hot Start Master Mix, 4 µL of 1:4 dilution of forward fragment, 4 µL of 1:4 reverse fragment, 2 µL 5’ N1-For at 4.5 µM, 2µL 3’ N1-Rev at 4.5µM and 3µL water. The PCR cycling conditions were the same as described above, but using 20 cycles instead of 7. The joined products were purified, diluted to 3 ng/µL and used as a template for a second round of mutagenesis and joining PCR reactions. The joined products from the second round were then used for ligation.

*Gibson assembly for preparing plasmid library*

The NA PCR library was utilised to generate an NA plasmid library via Gibson assembly and high efficiency transformation. The pHW2000 plasmid was linearized by BsmBI enzyme digestion and dephosphorylated with Thermosensitive Alkaline Phosphatase (Promega, USA). The assembly for the joined products were set up in duplicates, where 100 ng of purified vector was incubated with 3-fold excess of joined PCR product and 10 µL of 2x Gibson Assembly Master Mix (New England Biolabs, USA), in a total reaction volume of 20 µL. The reactions were incubated at 50ºC for 60 minutes, pooled together in a total volume of 40 µL and purified with AMPure XP Beads (bead to sample ratio = 1.8) (Beckman-Coulter, USA).

The purified assembled products were transformed into high efficiency electro-competent MegaX DH10B™ T1R Electrocomp™ cells (Thermo Fisher Scientific, Australia). Briefly, 2.5 µL of assembled products were mixed with 20 µL of bacterial cells and transferred to a 0.1 cm chilled cuvette and electroporation was performed at 2 kV. The bacterial cells were immediately resuspended in 250 µL SOC media and incubated for 1 hour at 37ºC, shaking at 220 rpm, and then 200 µL was plated onto agar plates. A 1/2000 dilution of the transformed bacterial cells were also plated out for colony counting. At least five independent transformations were carried out for each preparation of the assembled products. The following day there were between 200,000 to 400,000 colonies on each agar plate, as determined by colony counting from the diluted agar plates. After pooling colonies from the undiluted agar plates (2,000,000 x 5 = 10^6^ colonies total), they were cultured for 4 hours at 37ºC and then maxi-preps were performed using the EndoFree® Plasmid Maxi kit (Qiagen, Germany).

To gain insight regarding the distribution of codon mutations in the plasmid libraries, a total of 54 clones were picked from the diluted agar plates (with equivalent numbers of clones taken from the three replicates), and Sanger sequencing was performed. The distribution of codon mutations in the Sanger-sequenced clones were further analysed using a custom python script (<https://github.com/jbloomlab/SangerMutantLibraryAnalysis>).The three plasmid libraries (i, ii, and ii) were deep sequenced to determine if all possible substitutions had been comprehensively sampled.

*Reverse genetics for generation of virus library*

Each NA plasmid library was used for reverse genetics to generate a virus library (Figure 1C) [59, 61, 63]. Co-cultures of 293T and MDCK-SIAT1-TMPRRS2 [64] cells in 6-well plates were transfected with pHW2000 containing seven genes from A(H1N1)pdm09 virus A/South Australia/16/2017, and with different NA plasmid preparations containing H275Y-NA from the same virus. Overall, four viral rescues were performed: three with NA plasmid library replicates and one control rescue with the non-library H275Y-NA plasmid. Each rescue was carried out in six replicates, and the supernatants from the different wells were pooled together to create the final virus library.

Prior to transfection, 4x10^5^ 293T cell and 0.5x10^5^ MDCK-SIAT-TMPRSS2 cells were seeded in 6-well plates and grown for 24 hours in GIBCO Opti-Mem® + GlutaMax (Life technologies, US) supplemented with 5% (v/v) foetal bovine serum (JRH Biosciences, US), 2 µM L-glutamine (SAFC Biosciences, US), 200 U/mL penicillin (Sigma-Aldrich, US), 200 µg/mL streptomycin (Sigma-Aldrich, US ), 0.02M HEPES(SAFC Biosciences) and 2 mg/L Amphotericin B Fungizone (Sigma‑Aldrich, US). Viral rescues were performed with equal amounts of the NA plasmid library and the seven internal genes of the A/South Australia/16/2017 virus (HA, NP, NS, M, PA, PB1, PB2) in the pHW2000 plasmid. Growth media was removed from each well and transfected with a total of 1µg of plasmid pre-mixed with 1 ml GIBCO Opti-Mem® + GlutaMax and 18 µL FuGENE® HD transfection reagent (Promega, US). Co-cultures were incubated for 6 hours at 37°C, 5% CO_2_ with the transfection mixture, which was then replaced with 1 ml of GIBCO Opti-Mem® + GlutaMax (Life technologies, US) supplemented with 200 U/mL penicillin (Sigma-Aldrich, US), 200 µg/mL streptomycin (Sigma-Aldrich, US ) and 0.02 M HEPES(SAFC Biosciences). After another 24 hour incubation, 1ml of GIBCO Opti-Mem® + GlutaMax (Life technologies, US) supplemented with 4 µg/mL TPCK-trypsin (SAFC Biosciences, US), 200 U/mL penicillin (Sigma-Aldrich, US), 200 µg/mL streptomycin (Sigma-Aldrich, US ) and 0.02M HEPES( SAFC Biosciences) was added to each well. After a further 48 hour incubation, the supernatants in each well were tested for presence of virus using a standard hemagglutination (HA) test [65]. If HA titre was detected, the supernatant from the replicates in each rescue were pooled, centrifuged at 2000 rpm to remove cell debris and stored at -80 ºC. Titres of infectious virus in the virus library preparation were determined using a TCID_50_ assay [66].
