## Supplementary Figures for "Predicting potentially permissive substitutions that improve the fitness of A(H1N1)pdm09 viruses bearing the H275Y NA substitution"

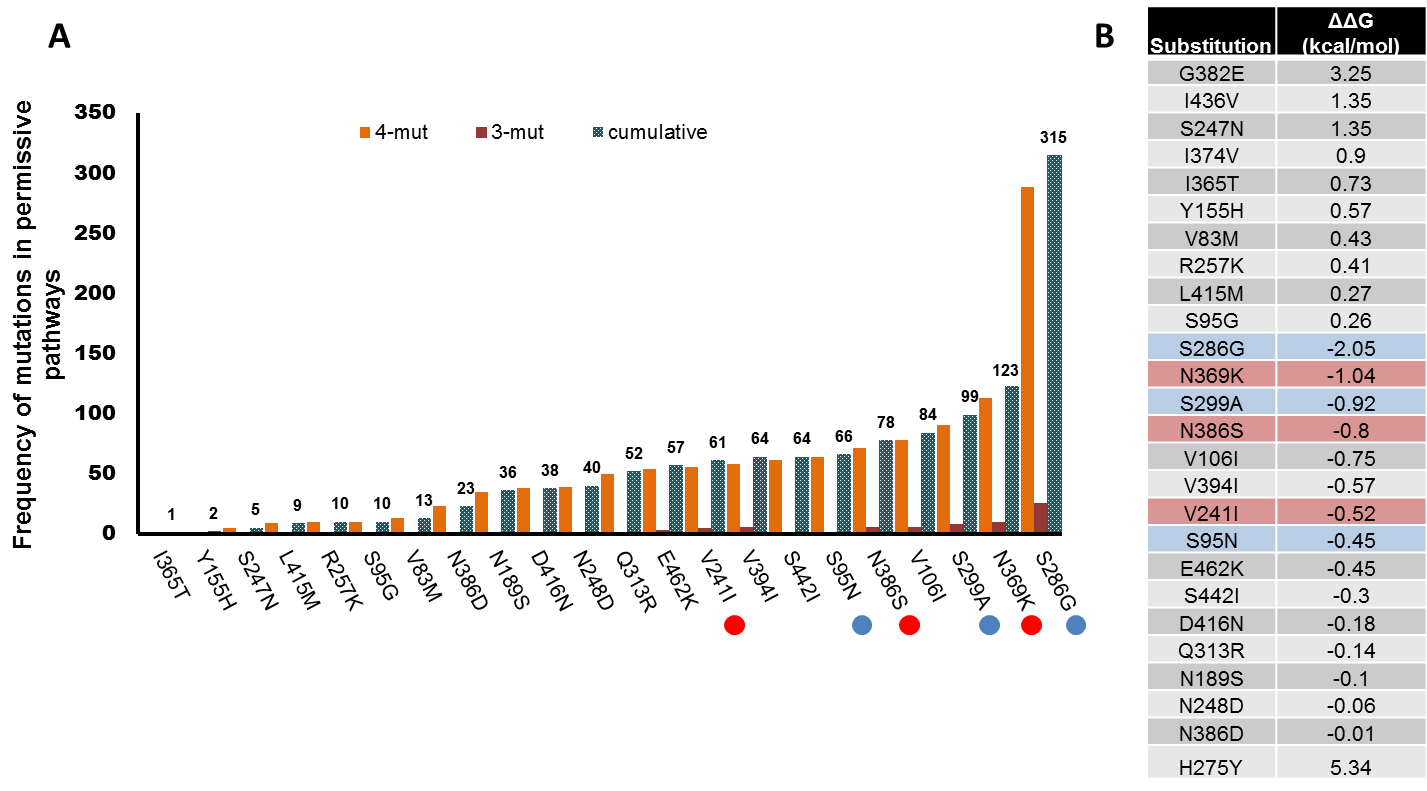

**Figure S1: Frequency and protein stability of substitution sets in permissive pathways identified by computational analyses of the viral N1 NA, (A)** The frequency of the 25 substitutions in the 3 and 4 substitution sets in the permissive pathways. **(B)** The *in silico* effect of each substitution on protein stability was calculated by FoldX, where a negative number represents increased protein stability and a positive number represents reduced stability. For comparison, the effect of H275Y in reducing protein stability was calculated and was shown to reduce in silico protein stability more substantially than any other substitution. The candidate substitutions S286G, S299A and S95N that were selected for experimental analysis are indicated by blue dots (Panel A) or highlighted in blue (Panel B). The permissive substitutions previously evaluated by Butler *et al*.[1] are indicated by red dots (Panel A) or highlighted in red (Panel B). Of note the V106I substitutions was not chosen for further evaluation as it has been replaced with the S200N substitution in currently circulating strains.

**Table S1: The nucleotide diversity of the NA gene in viruses from ferret nasal wash samples**

| **Ferret** | **Replicate No** | **π** | **πN/πS** |
| --- | --- | --- | --- |
| Experimentally infected | Replicate 1 | 0.0018 | 0.6261 |
|  | Replicate 2 | 0.0012 | 0.6046 |
|  | Replicate 3 | 0.0018 | 0.5296 |
|  | Replicate 4 | 0.0017 | 0.5299 |
|  | Control | NA | NA |
| Direct Contact 1 | Replicate 1 | 0.0006 | 0.6372 |
|  | Replicate 2 | 0.0004 | 0.8021 |
|  | Replicate 3 | 0.0011 | 0.8182 |
|  | Replicate 4 | 0.0023 | 0.1975 |
|  | Control | NA | NA |
| Direct Contact 2 | Replicate 1 | 0.0003 | 1.2227 |
|  | Replicate 2 | 0.0003 | 0.3404 |
|  | Replicate 3 | 0.0008 | 0.3786 |
|  | Replicate 4 | 0.0011 | 0.2736 |
| Aerosol Contact | Replicate 1 | 0.0003 | 0.5640 |
|  | Replicate 2 | 0.0002 | 0.9984 |
|  | Replicate 3 | 0.0002 | 0.6804 |
|  | Replicate4 | 0.0009 | 0.0624 |

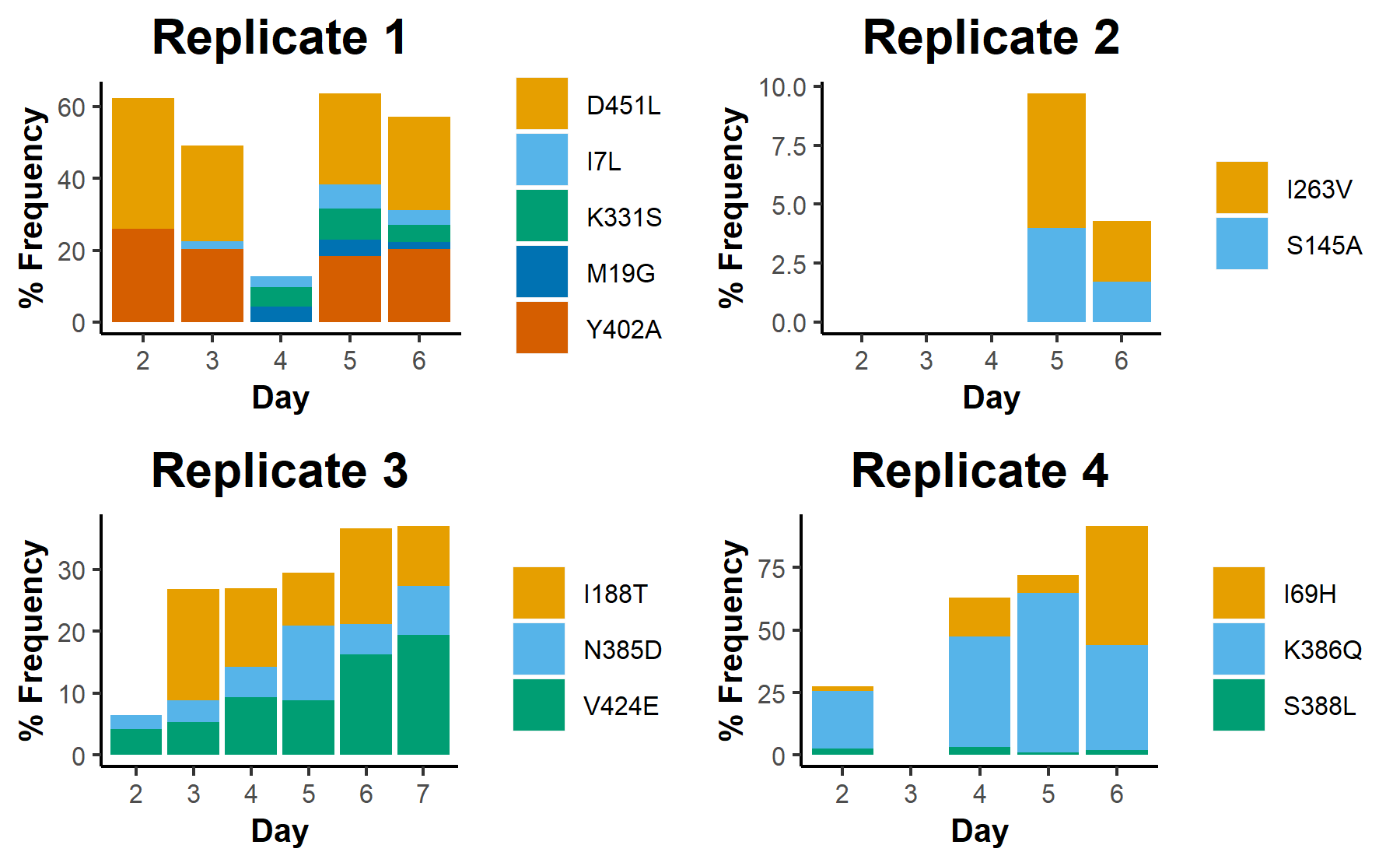

**Figure S2: Changes in frequency of variants across experimental days in Direct Contact 1 animals.** Nasal wash samples across the different experimental days were used to extract viral RNA which was then sequenced on the Illumina platform, and aligned using Bowtie2. The frequency of variants was calculated by Varscan. Most direct contact 1 animals stopped shedding infectious virus after day 6/7 and due to poor RNA quality no data is available for Day 3 in replicate 4, and Day 2,3, or 4 for Replicate 2.

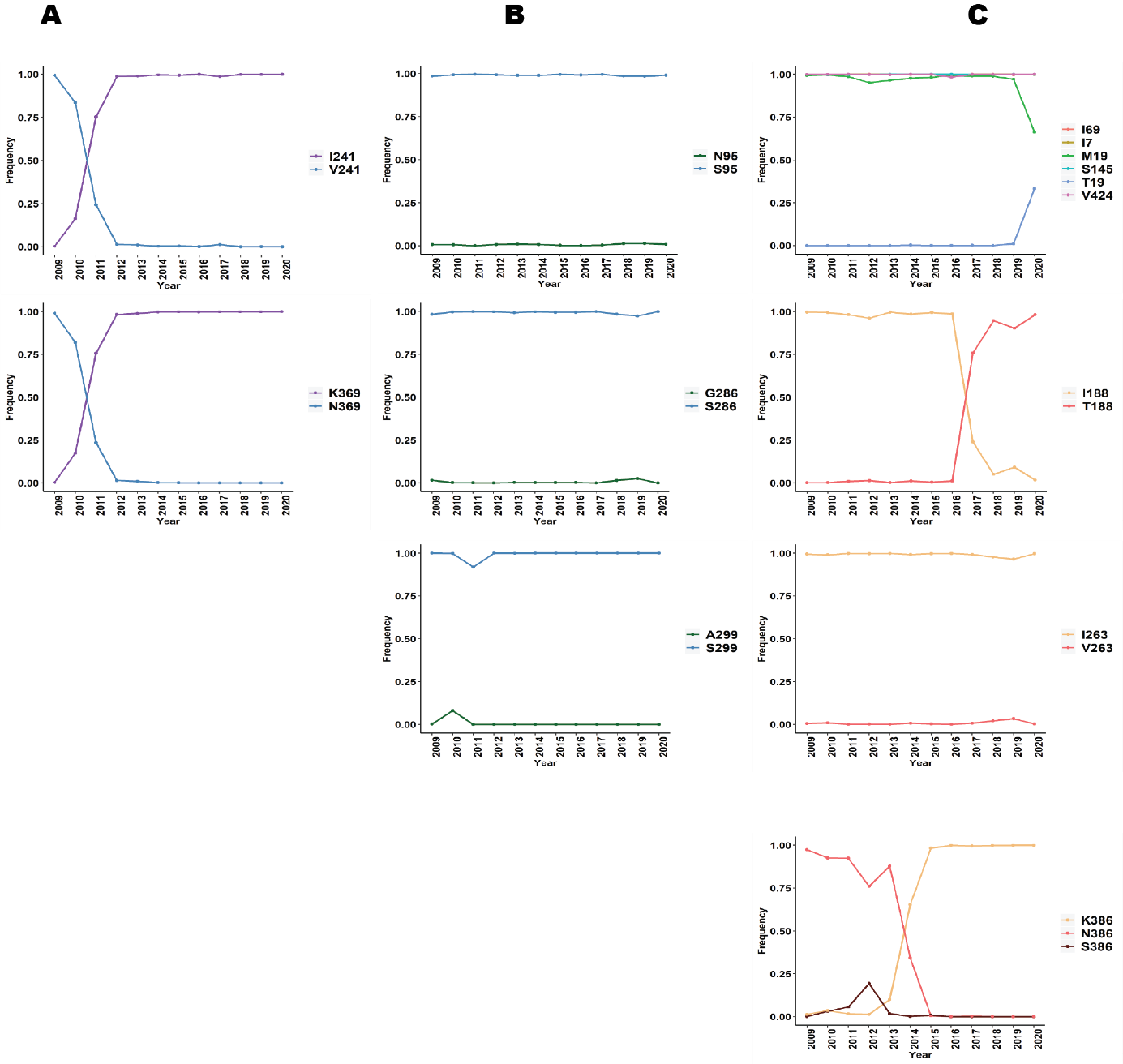

**Figure S3: Frequency of previously identified and novel candidate permissive substitutions in circulating N1 NA sequences of A(H1N1)pdm09 viruses.** All N1 NA protein sequences submitted to the GISAID sequence database since 2009 were downloaded and cleaned to remove short sequences and sequences with duplicate designations. The 34510 remaining sequences were then aligned using MAFFT and frequency of substitution at each position was calculated using a custom R script. **A)** Frequency of previously identified V241I and N369K permissive substitutions. **B)** Frequency of candidate substitutions proposed by computational analysis and **C)** Frequency of candidate substitutions proposed by experimental analysis.

1.	Butler, J., et al., *Estimating the fitness advantage conferred by permissive neuraminidase mutations in recent oseltamivir-resistant A(H1N1)pdm09 influenza viruses.* PLoS Pathog, 2014. **10**(4).
